# Platform-dependent effects of genetic variants on plasma APOL1

**DOI:** 10.1101/2025.01.30.635763

**Authors:** Qingbo S. Wang, Jinguo Huang, Leanne Chan, Nicole Haste, Niclas Olsson, Aleksandr Gaun, Fiona McAllister, Deepthi Madhireddy, Amos Baruch, Katie M. Cardone, Rachit Kumar, Marylyn Ritchie, Katalin Susztak, Eugene Melamud, Anastasia Baryshnikova

**Affiliations:** Calico Life Sciences LLC, South San Francisco, CA, USA; Flagship Pioneering, Cambridge, MA, USA; Department of Genetics and Institute for Biomedical Informatics, University of Pennsylvania Perelman School of Medicine, Philadelphia, PA, USA; Renal Electrolyte and Hypertension Division, Department of Genetics, Penn/CHOP Kidney Innovation Center, Institute for Diabetes Obesity and Metabolism, University of Pennsylvania, Philadelphia, PA, USA

## Abstract

Mutations in apolipoprotein L1 (APOL1) are strongly associated with protection against parasitic infections and increased risk of kidney disease in individuals of African ancestry. To better understand the mechanisms underlying APOL1-related pathologies, we examined genetic drivers of circulating APOL1 in individuals of African and European ancestry from four independent cohorts (UK Biobank, AASK, deCODE, and Health ABC) using three proteomic technologies (Olink, SomaLogic, and mass spectrometry). We found that disease-associated *APOL1* G1 and G2 variants are strong *cis*-pQTLs for plasma APOL1 measured by Olink and SomaLogic, but not mass spectrometry. Critically, the direction of variant effects differed between proteomic platforms, being positive with Olink and negative with SomaLogic. We identified an additional *APOL1* missense variant (rs2239785), common in Europeans, exhibiting the same platform-dependent directional discrepancy. Furthermore, variants in the kallikrein-kinin system (KKS), involving *KLKB1*, *F12*, and *KNG1*, and their genetic interactions showed strong *trans*-pQTL effects on APOL1 measured by Olink, but not SomaLogic. These platform-dependent discrepancies raise the possibility that both intrinsic *APOL1* mutations and extrinsic KKS activity induce conformational changes in the APOL1 protein that are differentially recognized by the proteomic platforms.

## Introduction

Chronic kidney disease (CKD) is a complex condition characterized by gradual loss of kidney function with multiple genetic and environmental influences (Obrador *et al*., 2017). Individuals of West African descent exhibit a disproportionately high risk of developing CKD, leading to increased rates of kidney failure and mortality (Ferguson, 1991). This ancestry-based disparity is largely attributed to coding variants in the *APOL1* gene, termed G1 and G2, which are common in African populations but virtually absent elsewhere (**Table 1**) (Genovese *et al*., 2010; Tzur *et al*., 2010). The G1/G2 variants have been linked to markedly increased odds ratio (OR) for several renal pathologies, including hypertension-associated end-stage renal disease (H-ESRD; OR = 7–10), focal segmental glomerulonephrosis (FSGS; OR = 10–17) and HIV-associated nephropathy (HIVAN; OR = 29–89) (Friedman and Pollak, 2020). Furthermore, mouse models with transgenic expression of these variants in podocytes develop features mirroring the human disease at functional, structural, and molecular levels (Beckerman *et al*., 2017).

**Table 1.** Key variants associated with plasma APOL1 levels.

| RSID | Name | Variant ID<br>(GRCh38) | Gene | Effect | Alternative allele<br>frequency<br>(gnomad) |  |
| --- | --- | --- | --- | --- | --- | --- |
|  |  |  |  |  | EUR | AFR |
| rs73885319 | G1G | 22:36265860:A:G | *APOL1* | S342G | $2.6 \times 10^{-4}$ | 0.22 |
| rs60910145 | G1M | 22:36265988:T:G | *APOL1* | I384M | $5.3 \times 10^{-5}$ | 0.23 |
| rs1317778148<br>rs71785313 | G2 | 22:36265995:AATA<br>ATT:A | *APOL1* | N388_Y<br>389del | $4.6 \times 10^{-5}$ | 0.14 |
| rs73885316 | M1 | 22:36265628:C:A | *APOL1* | N264K | $1.4 \times 10^{-4}$ | 0.03 |
| rs2239785 | V1 | 22:36265284:G:A | *APOL1* | E150K | 0.81 | 0.35 |
| rs6000220 | V2 | 22:36253920:C:T | *APOL1* | 5'-UTR | 0.11 | 0.16 |
| rs148296684 | – | 22:36265308:C:T | *APOL1* | L158F | $1.4 \times 10^{-3}$ | $1.3 \times 10^{-4}$ |
| rs1801020 | – | 5:177409531:A:G | *F12* | 5'-UTR | 0.75 | 0.55 |
| rs3733402 | – | 4:186236880:G:A | *KLKB1* | S143N | 0.51 | 0.74 |

The *APOL1* gene encodes apolipoprotein L1, a primate-specific protein expressed in multiple tissues and circulating in plasma as a component of high-density lipoprotein (HDL) particles (Shiflett *et al*., 2005). Circulating APOL1 provides protection against African sleeping sickness by forming cytotoxic ion channels in the disease-causing parasite *Trypanosoma brucei*, leading to its destruction (Friedman and Pollak, 2020). The G1 and G2 variants in *APOL1*, resulting in two amino acid substitutions (referred to as G1M and G1G, in near perfect linkage disequilibrium) and a two amino acid deletion, respectively (**Table 1**), confer enhanced protection against a parasite subspecies, *T. brucei rhodesiense*, by preventing the binding of an APOL1-inhibiting factor (Friedman and Pollak, 2020). This enhanced protection likely provided a survival advantage to G1/G2 carriers in regions where the parasite is prevalent (Friedman and Pollak, 2011).

While the roles of APOL1 and its G1/G2 mutations in *Trypanosoma* resistance are well-established, the mechanisms by which G1/G2 increase CKD risk are less clear. A prevailing hypothesis suggests that, similar to its function in parasite defense, APOL1 acts as an ion channel in human podocytes. The G1/G2 variants are thought to dysregulate APOL1’s channel function, causing ionic imbalance and kidney cell death (Lan *et al*., 2014; Olabisi *et al*., 2016; Giovinazzo *et al*., 2020). This hypothesis is supported by promising results from preclinical and early clinical studies indicating that blocking APOL1 channels prevents podocyte injury, reduces proteinuria and preserves kidney function (Egbuna *et al*., 2023; Vasquez-Rios, De Cos and Campbell, 2023). While most research has focused on APOL1 expressed in the kidneys, the role of circulating APOL1 has been less extensively studied, likely due to early reports of no association between plasma APOL1 levels and kidney disease (Bruggeman *et al*., 2014; Kozlitina *et al*., 2016).

Recent advancements in proteome profiling methods have greatly expanded our capacity to investigate the contribution of circulating proteins to disease. In particular, high throughput proteomic methods combined with genetic analyses of large collections of human plasma samples have driven the discovery of disease-relevant protein quantitative trait loci (pQTLs), i.e. genetic variants associated with both disease risk and altered protein levels in the plasma (Zhang *et al*., 2022; Sun *et al*., 2023). Large-scale plasma pQTL studies are enabled primarily by two high throughput and high sensitivity technologies: the proximity extension assay (PEA) commercialized by Olink, which uses target-specific antibodies tagged with molecular barcodes (Fredriksson *et al*., 2002), and the DNA aptamer technology commercialized by SomaLogic, which relies on target-specific single-stranded oligonucleotides (Gold *et al*., 2010). While both technologies provide quantitative (though relative) and internally reproducible measurements of protein abundance, direct comparisons have revealed a wide range of correlations between the two platforms (Eldjarn *et al*., 2023).

In this study, we present a comprehensive analysis of APOL1 plasma pQTLs in individuals of African descent, comparing data from the UK Biobank, generated with Olink’s technology, and the African American Study of Kidney Disease and Hypertension (AASK), generated with SomaLogic’s technology. Our findings reveal a key difference: while the G1/G2 mutations have strong pQTL effects on both platforms, they are associated with increased APOL1 plasma levels when measured by Olink and decreased levels when measured by SomaLogic. By comparing pQTLs in European individuals across both platforms, we confirmed that this discrepancy is specific to protein-altering variants in the *APOL1* gene itself, rather than non-coding *cis*-regulatory variants. Furthermore, we find that genes in the kallikrein-kinin system (KKS; *KLKB1*, *F12*, and *KNG1*) have strong *trans*-pQTL effects on APOL1, but these effects are only observed in the Olink data. We hypothesize that these platform-specific effects may arise from differential affinity of Olink and SomaLogic reagents for conformational isoforms of APOL1, induced by its coding mutations and potentially by proteolytic cleavage via the KKS pathway.

## Results

### G1 and G2 mutations increase Olink-based estimates of plasma APOL1 levels (APOL1_Olink_) in UKBB

To explore a potential role of circulating APOL1 in pathology, we analyzed the association between the G1/G2 risk variants and plasma APOL1 levels in 828 individuals of African descent (AFR) from the UK Biobank (Sudlow *et al*., 2015). Due to concerns regarding the accuracy of imputation for the G1 variant (**Fig. S1**), we preferentially used exome sequencing data where available, resorting to imputed data in all other instances (Methods). As the UK Biobank study used Olink’s PEA technology for relative protein quantification (Fredriksson *et al*., 2002), we denote these measurements as APOL1_Olink_ to distinguish them from data obtained via other platforms.

The *APOL1* locus showed a strong association signal with plasma APOL1_Olink_ (p < 10^−100^; **Fig. 1A**), with multiple variants correlating with elevated protein levels. Statistical fine-mapping of the region provided strong evidence that the G1 and G2 mutations are the causal determinants of this association, with a posterior inclusion probability (PIP) exceeding 0.99 for G2 and 0.76 for G1 (sum of 0.44 for G1M and 0.32 for G1G; **Fig. 1A**). Due to the near-perfect linkage between G1G and G1M, we used G1M as a proxy for both variants in subsequent analyses. Notably, due to the linkage between G1 and G2 (Pearson R = –0.27), the effect of G1 on APOL1_Olink_ levels is only detected after adjusting for G2 (p = 0.23 and p = 1.3×10^−19^ before and after adjustment, respectively). Adjusting for both G1 and G2 abolished the association signals of all other variants (p < 10^−100^ and p > 5×10^−8^ before and after adjustment, respectively), confirming that G1 and G2 are the main *cis*-genetic determinants of plasma APOL1_Olink_ levels in the AFR population (**Fig. 1A**).

**Figure 1.**
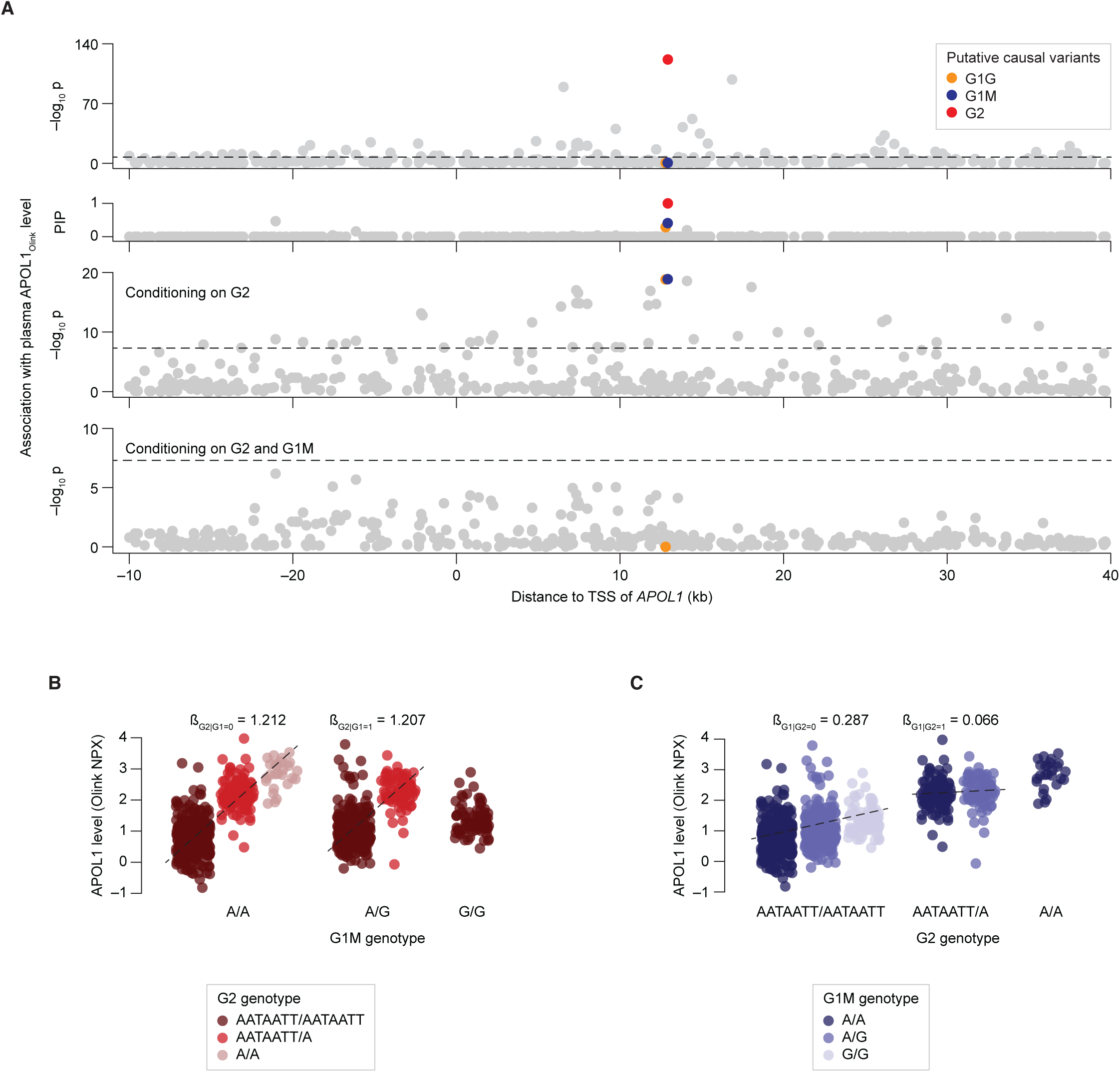
*APOL1* G1 and G2 variants are causal pQTLs for plasma APOL1_Olink_ levels. **(A)** Manhattan plot showing the association of *APOL1 cis*-variants with plasma APOL1_Olink_ levels in the UK Biobank population of African descent. Each dot represents a variant. The x-axis shows the variant’s chromosomal position relative to the transcriptional start site (TSS) of *APOL1*. The y-axis shows the –log_10_ of the association p-value (row 1), the statistical fine-mapping posterior inclusion probability (PIP; row 2), the –log_10_ of the association p-value after adjusting for the effect of the G2 variant (row 3) and the –log_10_ of the association p-value after adjusting for the effects of both the G2 and G1M variants (row 4). Putative causal variants identified through fine-mapping are highlighted. Horizontal dashed lines represent the genome-wide significance threshold (p = 5×10_–8_). **(B)** Scatter plot showing the effect of the G1M variant on plasma APOL1 levels, stratified by G2 genotype. **(C)** Scatter plot showing the effect of the G2 variant on plasma APOL1_Olink_ levels, stratified by the G1M genotype. **(B–C)** Each dot represents an individual. Within each genotype group, dots were jittered along the x-axis for visualization purposes.

Our analysis of G1/G2 effect on plasma APOL1_Olink_ levels favored an additive model of inheritance (**Fig. 1B–C**). Specifically, each copy of the G1 or G2 risk allele contributed to increasing APOL1_Olink_, such that heterozygous individuals exhibited intermediate APOL1_Olink_ levels relative to those with zero or two copies of the variant (**Fig. 1B–C**). Notably, the effect of G2 in the absence of G1 (β_G2|G1=0_ = 1.212) was considerably stronger than the effect of G1 without G2 (β_G1|G2=0_ = 0.287; **Fig. 1B–C; Fig. S2A**). The presence of G2 appeared to further attenuate the effect of G1 (β_G1|G2=1_ = 0.066; **Fig. 1C**), suggesting that G1 and G2 share a genetic interaction (p = 0.0094 for difference between β_G1|G2=0_ and β_G1|G2=1_; Methods). Similarly, G2 showed a moderate but significant genetic interaction with the APOL1 M1 variant (rs73885316, also known as p.N264K; **Table 1**): while M1 did not exhibit an effect on its own, it attenuated the effect of G2 (β_G2|(G1=0&M1>0)_ = 0.985; p = 0.032 for difference between β_G2|(G1=0&M1=0)_ and ^β^(G2|G1=0&M1>0)^; **Fig. S2F**).^

### The effects of G1 and G2 mutations on plasma APOL1 levels are platform-specific

We sought to replicate the observed pQTL effects of G1/G2 mutations on plasma APOL1_Olink_ in an independent cohort of similar size and genetic ancestry: the African American Study of Kidney Disease and Hypertension (AASK) (Appel *et al*., 2003). This study includes 461 African American participants with plasma APOL1 levels measured using SomaLogic’s aptamer technology (Surapaneni *et al*., 2022). We denote these measurements as APOL1_SomaLogic_. Analysis of pQTL summary statistics from AASK revealed a significant association between the G2 mutation and APOL1_SomaLogic_ levels (p = 3.2×10^−8^). However, in contrast to our UK Biobank findings, this mutation was associated with an apparent decrease in APOL1 concentration (β = –0.45; **Fig. 2A**; **Table 2**). The direction of the effect of G1 was consistent with G2 (β = –0.10; **Fig. 2A**; **Table 2**), although it did not reach genome-wide significance likely due to linkage with G2.

**Figure 2.**
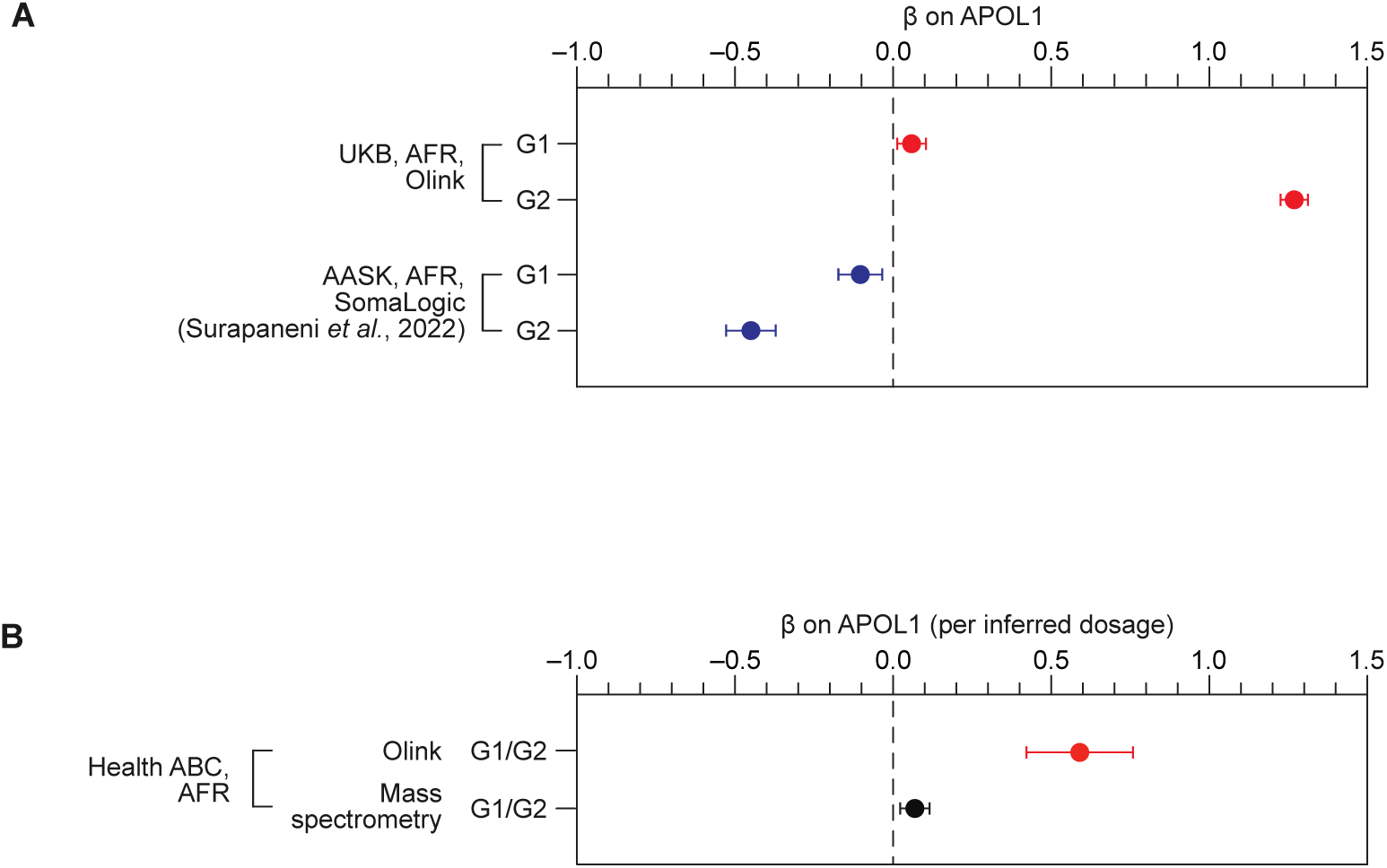
*APOL1* G1 and G2 variants show opposite effects on plasma APOL1 levels measured by Olink and SomaLogic platforms. **(A)** Association of *APOL1* G1 and G2 variants with plasma APOL1 levels in individuals of African ancestry from the UK Biobank (Olink platform) and AASK study (SomaLogic platform). Effect sizes are reported as posterior β (UK Biobank) and marginal β (AASK study). Error bars indicate standard errors of the mean. (B) Association of *APOL1* G1/G2 variants with plasma APOL1 levels measured by Olink’s platform and mass spectrometry in individuals of African ancestry from Health ABC. Error bars indicate standard errors of the mean.

**Table 2.** Association of *APOL1* coding variants with plasma APOL1 levels across multiple datasets and platforms. Values represent β coefficients of the specified variant allele. In the UKB AFR dataset, posterior effect sizes derived from statistical fine-mapping are shown to adress linkage disequilibrium (LD) between V1 and G1/G2. In the AASK dataset, the proxy variant rs12106505 was used for G2 (Bajaj *et al*., 2020) as G2 was absent from the summary statistics. In the Health ABC dataset, G1 and G2 are indistinguishable **(Methods**). Dash lines (“–”) indicate that data were not available; “n.s.” indicates that the association was not significant.

| Dataset | Ancestral group | Platform | Sample size | *APOL1* coding variant ($\beta$) | | |
| --- | --- | --- | --- | --- | --- | --- |
|  |  |  |  | G1 | G2 | V1 (G allele) |
| UKB | AFR | Olink | 828 | 0.28 | 1.33 | 0.18 |
| UKB | EUR | Olink | 38,442 | – | – | 0.32 |
| AASK | AFR | SomaLogic | 461 | n.s. | -0.44<br>(rs12106505) | -0.33 |
| deCODE | EUR | SomaLogic | 35,559 | – | – | -0.38 |
| Health ABC | AFR | Olink | 194 | 0.59<br>(G1/G2) | 0.59<br>(G1/G2) | 0.27 |
| Health ABC | EUR | Olink | 366 | – | – | 0.14 |
| Health ABC | AFR | MS | 390 | n.s. | n.s. | n.s. |
| Health ABC | EUR | MS | 704 | – | – | n.s. |

To address the conflicting pQTL effects observed between the UK biobank (Olink) and AASK (SomaLogic), we sought further evidence from a third independent cohort: the Health, Aging and Body Composition (Health ABC) study (Newman *et al*., 2023). We used Olink’s technology and mass spectrometry to measure plasma APOL1 levels in 560 and 1,094 Health ABC participants (including 194 and 390 individuals of African descent), respectively (Methods). As genotype data were not directly available, we inferred G1/G2 mutation status using peptide sequence analysis (Methods; **Fig. S3A**). This inference was based on the principle that the abundance of the APOL1 LNILNNNYK peptide, which spans both G1M and G2 variants, is expected to correlate with the number of reference allele copies at both loci and, consequently, can be used for predicting G1/G2 status (**Fig. S3A**).

Following genotype inference, our analysis of Health ABC data confirmed and further extended the platform-dependent G1/G2 effects (**Table 2**). Despite a significant positive correlation between APOL1_Olink_ and APOL1_MS_ levels (R = 0.39, p = 4.2×10^−20^; **Fig. S3C**), the presence of G1/G2 mutations was associated with increased APOL1_Olink_ levels (β = 0.59, p = 9.1×10^−4^), but showed no significant effect on APOL1_MS_ levels (β = 0.069, p = 0.16; p = 3.5×10^−3^ for heterogeneity of effect size between Olink and mass spectrometry measurements; **Fig. 2B, S3D**). The loss of pQTL effect with mass spectrometry cannot be attributed to decreased statistical power, given that the sample with mass spectrometry data (n = 390) is larger than the sample with Olink data (n = 194) and has similar population characteristics (**Table S1**).

These findings collectively suggest that, while G1/G2 mutations do not impact total APOL1 levels (as measured by mass spectrometry), they may affect Olink- and SomaLogic-based quantification, resulting in an apparent anti-correlation of pQTL effects. To investigate this observation further, we analyzed plasma APOL1 levels in European populations, which carry a different set of genetic variants relative to populations with African ancestry and thus may provide further insight into platform-dependent pQTL discrepancies.

### European populations exhibit platform-specific *cis-* and *trans*-pQTLs for APOL1

We conducted a genome-wide pQTL analysis of plasma APOL1 levels in UK Biobank participants with European ancestry (n = 38,442), measured using the Olink platform, and compared the results to pQTL summary statistics from the deCODE study (n = 35,559 Icelanders), which used SomaLogic’s technology (Ferkingstad *et al*., 2021). Importantly, a previous analysis directly comparing Olink and SomaLogic measurements in 1,500 Icelandic plasma samples demonstrated a high correlation between the two platforms for overall APOL1 levels (Spearman ⍴ = 0.69) (Eldjarn *et al*., 2023).

Our comparison identified a shared *cis*-pQTL signal at the *APOL1* locus (chromosome 22) and a shared *trans*-pQTL signal at the *HPR* locus (chromosome 16), as well as three Olink-specific *trans*-pQTLs at the *KNG1*, *KLKB1* and *F12* loci (chromosomes 3, 4 and 5, respectively; **Fig. 3A**). The effects of the shared *cis*-pQTLs exhibited a strong negative correlation between Olink and SomaLogic measurements (**Fig. 3B**), whereas the effects of shared *trans*-pQTLs were positively correlated (**Fig. 3C**). To investigate potential mechanisms underlying *cis-* and *trans-*pQTL platform discrepancies, we examined each pQTL signal in detail.

**Figure 3.**
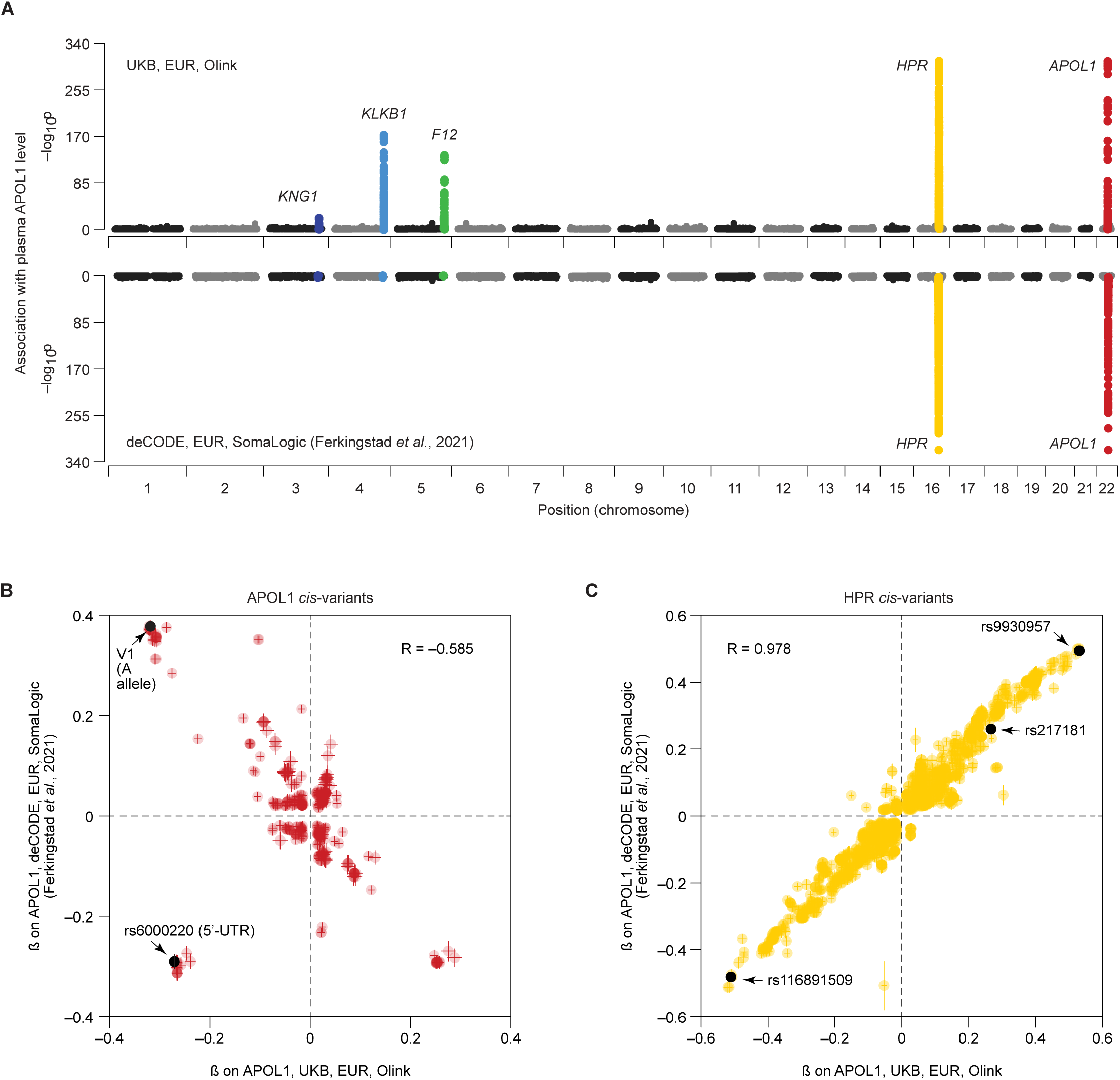
Platform-specific pQTLs influence plasma APOL1 levels through both *cis* and *trans* mechanisms. **(A)** Miami plot showing the genome-wide variant associations with plasma APOL1 levels in the UK Biobank (top, Olink) and deCODE (bottom, SomaLogic) populations of European ancestry. Each dot represents a variant. The x-axis shows chromosomal position. The y-axis shows the –log10 of the association p-value. The top five strongest associations are labeled with the names of the corresponding genes. **(B–C)** Scatter plots comparing the effect sizes (marginal β) of *cis-APOL1* (B) and *cis-HPR* (C) variants on plasma APOL1 levels in the UK Biobank (x-axis) and deCODE (y-axis).

### An APOL1 missense variant common in Europeans shows functional similarity to G1/G2

The APOL1 pQTL signal on chromosome 22 localized to the *APOL1* gene region (**Fig. 4A**), and was strong (p < 10^−320^) despite the virtual absence of G1/G2 mutations in European populations. Statistical fine-mapping of this region in the UK Biobank dataset identified three variants as potentially causal for the association: a common missense variant (rs2239785, hereafter referred to as V1; PIP = 0.75; **Table 1**), a common 5’-UTR variant (rs6000220, hereafter referred to as V2; PIP > 0.99; **Table 1**) and a rare missense variant (rs148296684; PIP > 0.99; **Table 1**). Due to its low frequency (MAF < 0.001, explaining less than 0.2% of the total variance in APOL1_Olink_), we could not further characterize the rare missense variant. Adjusting for the two common variants substantially diminished the association signal of all other variants in the region (p < 10^−320^ and p > 10^−20^ before and after adjustment, respectively; **Fig. 4A**).

**Figure 4.**
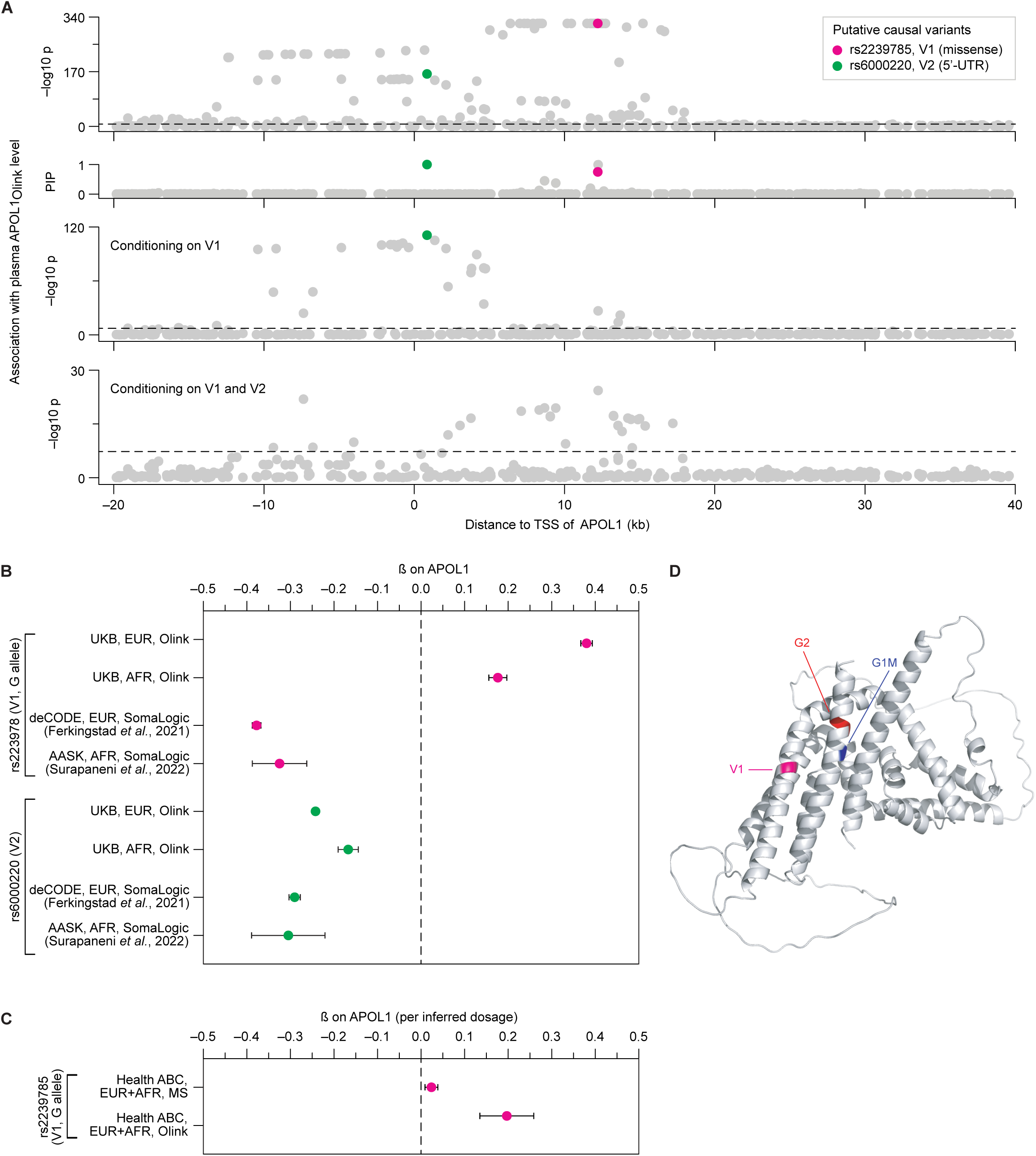
An *APOL1* missense variant common in European populations shows functional similarity to G1/G2. **(A)** Manhattan plot showing the association and statistical fine-mapping of *APOL1* cis-variants with plasma APOL1 levels in the UK Biobank population of European ancestry. Each dot represents a variant. The x-axis shows the variant’s chromosomal position relative to the transcriptional start site (TSS) of *APOL1*. The y-axis shows the –log10 of the association p-value (row 1), the fine-mapping posterior inclusion probability (PIP; row 2), the –log10 of the association p-value after adjusting for the effect of rs2239875 (V1; row 3) and the –log10 of the association p-value after adjusting for the effects for both V1 and rs6000220 (V2; row 4). Putative causal variants identified through fine-mapping are highlighted. Horizontal dashed lines represent the genome-wide significance threshold (p = 5×10^−8^). **(B)** Effects of the *APOL1* rs2239875 (V1) and rs6000220 variants on plasma APOL1_Olink_ and APOL1_SomaLogic_ levels in individuals of European ancestry from the UK Biobank and deCODE. Effect sizes are reported as posterior beta (UK Biobank, Olink) and marginal beta (deCODE, SomaLogic). Error bars indicate the standard error of the mean. **(C)** Association of the *APOL1* V1 variant with plasma APOL1 levels measured by Olink’s platform and mass spectrometry in individuals from Health ABC. Error bars indicate standard errors of the mean. **(D)** Predicted APOL1 protein structure (AlphaFold 2.0, visualized by PyMOL). Positions of the G1, G2 and V1 variants are highlighted.

Comparing the effects of the missense variant (V1) and the 5’-UTR variant (V2) between the UK Biobank (Olink) and deCODE (SomaLogic) revealed a notable difference. While the V2 variant consistently showed a negative association with APOL1 in both datasets, the V1 variant exhibited opposite directional effects (**Fig. 3B, 4B**). This difference was also evident in the AFR populations from the UK Biobank and AASK, where both variants are common and independent from G1/G2 (**Fig. 4B**; **Table 2**). Specifically, the V1 G allele, which is more frequent in AFR than in EUR populations (**Table 1**), was associated with higher APOL1 levels as measured by Olink (posterior β = 0.38 in EUR and 0.18 in AFR; Methods) and lower levels as measured by SomaLogic (β = –0.38 in EUR and –0.33 in AFR), mirroring the pattern observed for the G1/G2 mutations, although with substantially smaller effect sizes (**Fig. 4B**; **Table 2**). Furthermore, consistent with the G1/G2 results, the effect of V1 was replicated in the Health ABC data measured by Olink and was significantly weaker when measured by mass spectrometry (p = 7.7×10^−3^ for heterogeneity of effect size; **Fig. 4C, S3D**). In Health ABC, V1 genotypes were inferred from peptide sequence analysis, using a methodology that differed slightly from the approach used for G1/G2 inference (Methods; **Fig. S3B**). These genotypes were then tested for association with APOL1 levels using a combined sample of EUR and AFR individuals to increase power (Methods).

Interestingly, despite being located ∼200 amino acids apart in the linear protein sequence, the V1 and G1/G2 mutations are in close spatial proximity within the predicted three-dimensional structure of APOL1, as determined by AlphaFold (Jumper *et al*., 2021) (**Fig. 4D**).

### HPR, an APOL1 binding partner, is a strong *trans*-pQTL for plasma APOL1 levels

The APOL1 pQTL signal on chromosome 16 localized to the *HPR* gene region (**Fig. 3A**). *HPR* encodes haptoglobin-related protein, an essential component of the HDL particles that carry APOL1 in the bloodstream (Vanhollebeke and Pays, 2010). HPR mediates the entry of APOL1-HDL complexes into *Trypanosoma* by interacting with a parasite cell surface receptor (Vanhollebeke and Pays, 2010). Consistent with this close functional relationship, plasma levels of HPR and APOL1, as measured by mass spectrometry in the Health ABC dataset, were highly correlated (R = 0.79, p = 1.2×10^−234^; **Fig. S4A**). In contrast, APOL1 showed a much weaker correlation with APOA1 (R = 0.16, p = 5.4×10^−8^), consistent with the fact that APOA1 is a component of most HDL particles and only 10% of these particles also carry APOL1 (Madhavan and O’Toole, 2014).

While statistical fine-mapping of the *HPR* region in the UK Biobank dataset identified three intronic variants with PIP > 0.9 (**Fig. S4B**), their functional interpretation remains challenging. However, the high correlation of variant effect sizes between the UK Biobank (Olink) and deCODE (SomaLogic) datasets (R = 0.98; **Fig. 3C**) strongly supports a consistent effect of HPR variants on plasma APOL1 levels, regardless of the proteomic platform.

Interestingly, previous studies have identified copy number variation (CNV) in *HPR*, unique to AFR populations and associated with *Trypanosoma* resistance (Hardwick *et al*., 2014). However, our fine-mapping analysis did not support an effect of *HPR* CNV on plasma APOL1_Olink_ levels (p = 1.3×10^−3^, PIP = 0; **Fig. S4C–D**).

### Olink-specific *trans*-pQTLs identify the kallikrein-kinin system as a potential novel regulator of plasma APOL1

In contrast to the *HPR trans*-pQTL effects, which were consistent across Olink and SomaLogic platforms, effects associated with the *KNG1*, *KLKB1* and *F12* loci were only detected by Olink (**Fig. 3A**). These three genes encode core components of the kallikrein-kinin system (KKS), a pathway regulating multiple physiological processes, including blood coagulation, inflammation, and blood pressure control (Wisniewski, Gangnus and Burckhardt, 2024) (**Fig. 5A**). Specifically, *KLKB1* and *F12* encode the serine protease precursors pre-kallikrein and factor XII, respectively, which undergo mutual cleavage to yield the active enzymes kallikrein and factor XIIa (Motta, Juliano and Chagas, 2023) (**Fig. 5A**). *KNG1* encodes high-molecular-weight kininogen (HMWK), which serves as a cofactor and substrate for kallikrein in several branches of the KKS cascade (Motta, Juliano and Chagas, 2023) (**Fig. 5A**).

**Figure 5.**
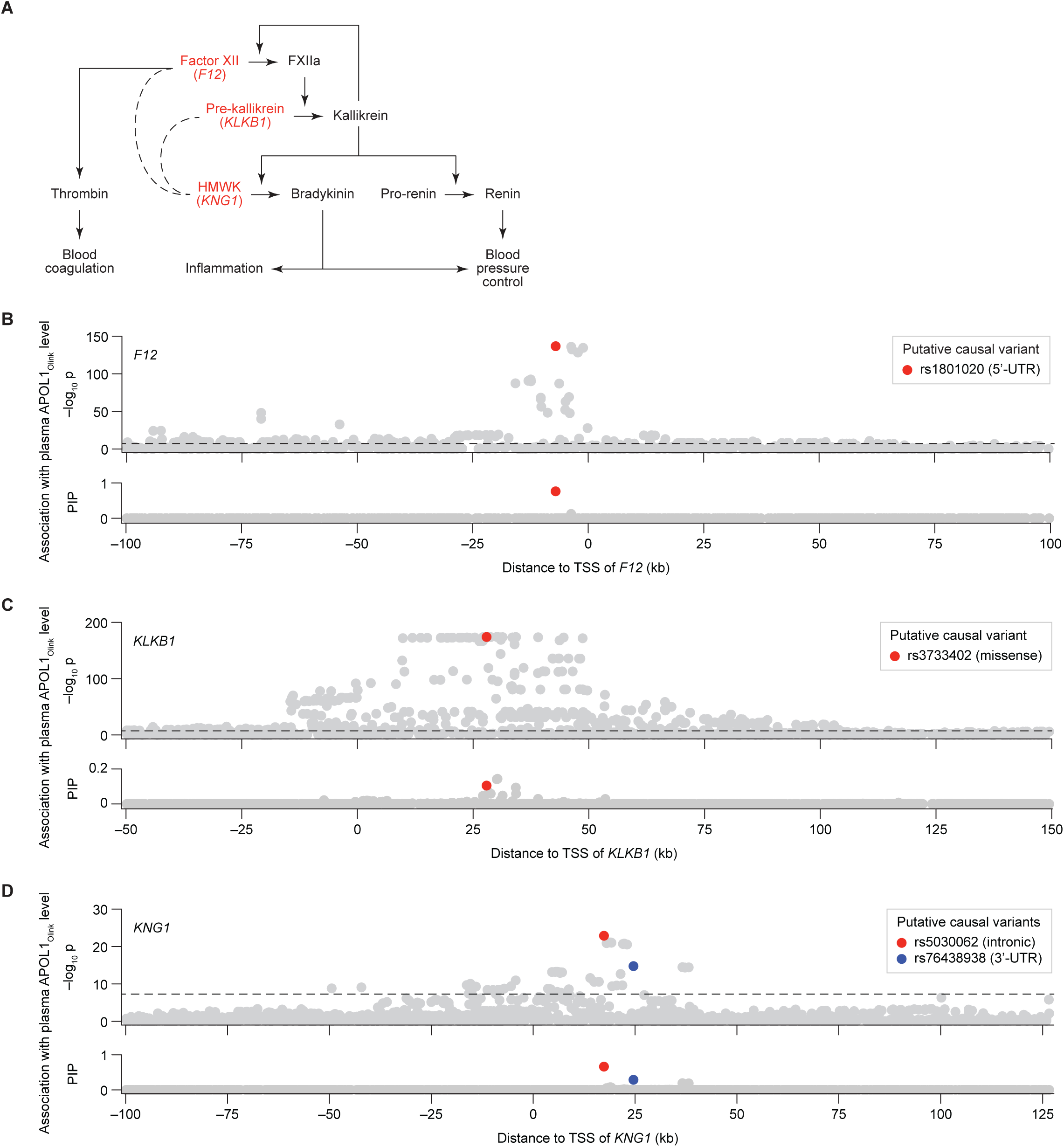
Variants in *F12*, *KLKB1* and *KNG1*, acting in the kallikrein-kinin pathway, are *trans*-pQTLs for plasma APOL1_Olink_ levels. **(A)** A schematic illustration of the kallikrein-kinin system, highlighting the relative influence of factor XII (FXII, encoded by *F12*), pre-kallikrein (encoded by *KLKB1*) and high molecular weight kininogen (HMWK, encoded by *KNG1*) on blood coagulation, inflammation and blood pressure control mechanisms. Arrows indicate enzymatic reactions and regulation. Dashed lines indicate co-factor/enzyme relationships. **(B–D)** Manhattan plots showing the association and statistical fine-mapping of *cis-F12* **(B)**, *cis-KLKB1* **(C)** and *cis-KNG1* **(D)** variants with plasma APOL1_Olink_ levels in the UK Biobank population of European ancestry. Each dot represents a variant. The x-axis show the variant’s chromosomal position relative to the transcriptional start site (TSS) of the indicated gene (*F12*, *KLKB1* or *KNG1*). The y-axis shows the –log_10_ of the association p-value (row 1) and the fine-mapping posterior inclusion probability (PIP; row 2). Putative causal variants identified through fine-mapping are highlighted. Horizontal dashed lines represent the genome-wide significance threshold (p = 5×10_–8_).

To understand how the KKS pathway might influence plasma APOL1_Olink_ levels, we performed statistical fine-mapping of the *KLKB1*, *F12,* and *KNG1* gene regions (**Fig. 5B–D**), and compared the effect of the prioritized variants on APOL1_Olink_ to their effect on plasma thrombin levels (**Fig. S5A–B**). Thrombin (factor II) is a blood coagulation protease acting downstream of factor XIIa and kallikrein (Chee, 2014) (**Fig. 5A**), and can be used as a biomarker of KKS pathway activity.

The association signal at the *F12* locus was largely attributed to a single 5’-UTR variant (rs1801020, PIP = 0.76; **Table 1**; **Fig. 5B**). The A allele at this variant introduces an alternative translation starting site for *F12*, leading to decreased production of functional factor XII protein (**Fig. S5A**) (Calafell *et al*., 2010) and lower plasma thrombin levels (Pietzner *et al*., 2020) (**Fig. S5B**).

The *KLKB1* locus displayed a more complex association pattern with multiple variants in high linkage disequilibrium (**Fig. 5C**). While the top two associated variants are intronic (rs139276089 and rs12509935, both with PIP = 0.14), we hypothesize that the third-ranking variant (rs3733402, PIP = 0.10), which is a missense variant, may be the true causal driver. Prior evidence indicates that the G allele at rs3733402 impairs the ability of kallikrein to bind and cleave HMWK, effectively acting as a partial loss-of-function (LOF) mutation (**Fig. S5A**) (Katsuda *et al*., 2007). Consistent with its predicted LOF effect, rs3733402-G has been reported to associate with lower plasma thrombin levels (Pietzner *et al*., 2020) (**Fig. S5B**).

The mechanistic interpretation of the *KNG1* association signal (**Fig. 5D**) is less straightforward, as the two putatively causal variants fall into an intronic region (rs5030062; PIP = 0.66; **Table 1**) and the 3’-UTR (rs76438938; PIP = 0.29; **Table 1**). Nevertheless, the A allele at rs5030062, similar to the non-functional alleles in *F12* and *KLKB1* described above, has also been reported to associate with lower plasma thrombin levels (Pietzner *et al*., 2020) (**Fig. S5B**).

Collectively, these observations suggest that genetic variation within the *F12*, *KLKB1*, and *KNG1* loci impacts both plasma APOL1_Olink_ levels and KKS pathway activity, as measured by plasma thrombin levels. Importantly, we observed a consistent directional relationship: alleles associated with decreased thrombin were also associated with decreased APOL1_Olink_, and vice versa (**Fig. S5A–B**). This correlation, which is independent of blood pressure-related factors (**Fig. S5B**), suggests that higher KKS pathway activity may lead, directly or indirectly, to elevated APOL1_Olink_ levels. Interestingly, despite comparable KKS variant allele frequencies between the Icelandic (deCODE) and UK populations (**Table S2**), these variants showed no association with APOL1_SomaLogic_ levels in the deCODE cohort (**Fig. 3A**).

### GxG analysis strengthens the association between the KKS pathway and plasma APOL1_Olink_

While standard pQTL analysis has proven effective in identifying individual variants associated with protein levels, it offers limited insight into their complex interplay. Genetic interaction (or GxG) analysis overcomes this single-variant limitation by investigating combined variant effects, allowing identification of synergy/antagonism and inference of functional relationships. Despite its widespread use in dissecting molecular pathways in model organisms (Baryshnikova *et al*., 2013), identifying GxG interactions affecting human quantitative traits has been challenging.

We used GxG analysis to investigate whether APOL1_Olink_ pQTLs levels and, by extension, their cognate genes (*APOL1* itself, *HPR*, *F12*, *KLKB1* and *KNG1;* **Fig. 3A**) act independently or in concert. Our analysis revealed strong evidence for GxG interactions between the KKS genes *KLKB1*, *F12* and *KNG1*, and, to a lesser extent, between *HPR* and *APOL1* (**Fig. 6A; Fig. S6**). Specifically, the interaction between *KLKB1* and *F12* variants (p < 2×10^−66^) was such that the association of each variant with plasma APOL1_Olink_ levels was nearly abolished when the other gene carried a non-functional allele (**Fig. 6B**). This true epistatic effect indicates that both *F12* and *KLKB1* must be functional for proper regulation of APOL1_Olink_, mirroring their interdependent roles in the KKS (**Fig. 5A**). Similarly, non-functional alleles in *F12* and *KLKB1* nearly eliminated the association of *KNG1* with plasma APOL1_Olink_ levels (p < 2×10^−8^ and p < 4×10^−10^ for GxG effect, respectively); however, the reciprocal influence, consistent with the intronic nature of *KNG1* alleles, was much weaker (**Fig. S6C–D**). In contrast, variants in *KLKB1* and *HPR* combined additively to influence APOL1_Olink_ levels (p < 0.47 for GxG effect; **Fig. 6C**), indicating that they act independently.

**Figure 6.**
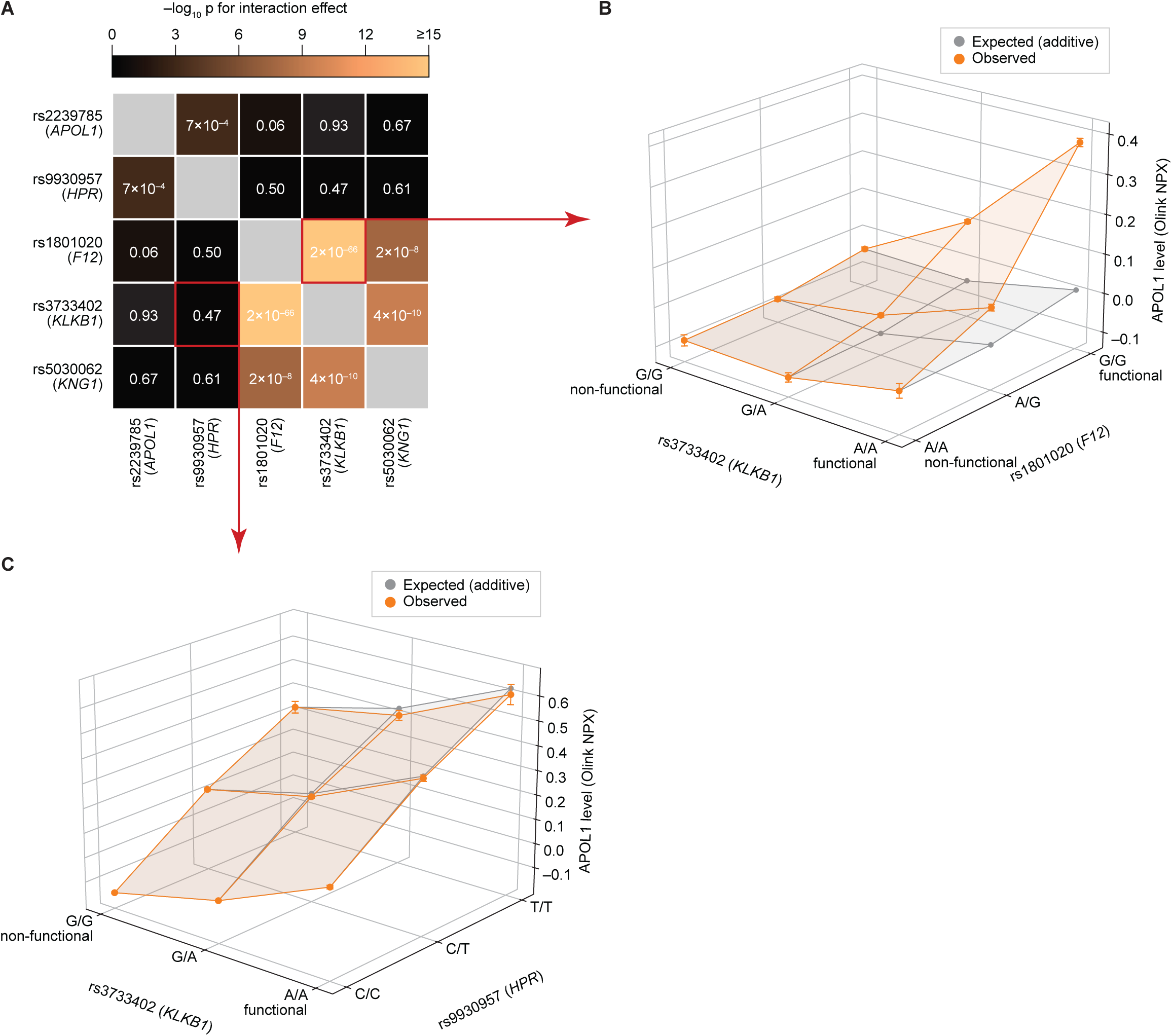
Variants in the kallikrein-kinin pathway genetically interact to influence plasma APOL1_Olink_ levels. **(A)** Heatmap showing the significance of pairwise genetic interactions between the top five putative causal pQTLs for plasma APOL1_Olink_ levels. Cell color indicates the –log_10_ of the interaction p-value (Methods). **(B–C)** Examples of an interacting **(B)** and a non-interacting **(C)** variant pair. In each plot, the x- and y-axes show variant genotypes, and the z-axis shows observed (orange) and expected, assuming additivity (grey), mean plasma APOL1_Olink_ levels for each genotype combination. Error bars indicate the standard error of the mean.

To further investigate the potential KKS-APOL1 link within the broader context of KKS pathway activity, we analyzed the impact of APOL1-associated KKS variants (rs3733402 for *KLKB1*, rs1801020 for *F12*, and rs5030062 for *KNG1*) on all measured plasma proteins in the UK Biobank. All three variants exhibited significant pleiotropy, influencing 133, 89, and 57 plasma proteins, respectively (p < 0.05 after Bonferroni correction). Despite this broad range of effects, their association with APOL1 consistently ranked among the top 5 individual and GxG effects (**Fig. 7A–B**). Other proteins ranking in the top 20 individual and GxG associations included established protease substrates and inhibitors, such as SERPINA5 (Protein C Inhibitor), SERPINI1 (Neuroserpin), ITGA6, ITIH4, and SPINT2 (**Fig. 7A–B**). Several of these inhibitors, notably SERPINA5 and ITIH4, have been reported to inhibit plasma kallikrein itself (Meijers *et al*., 1988; España, Berrettini and Griffin, 1989; Laurell and Stenflo, 1989; Pihl *et al*., 2021).

**Figure 7.**
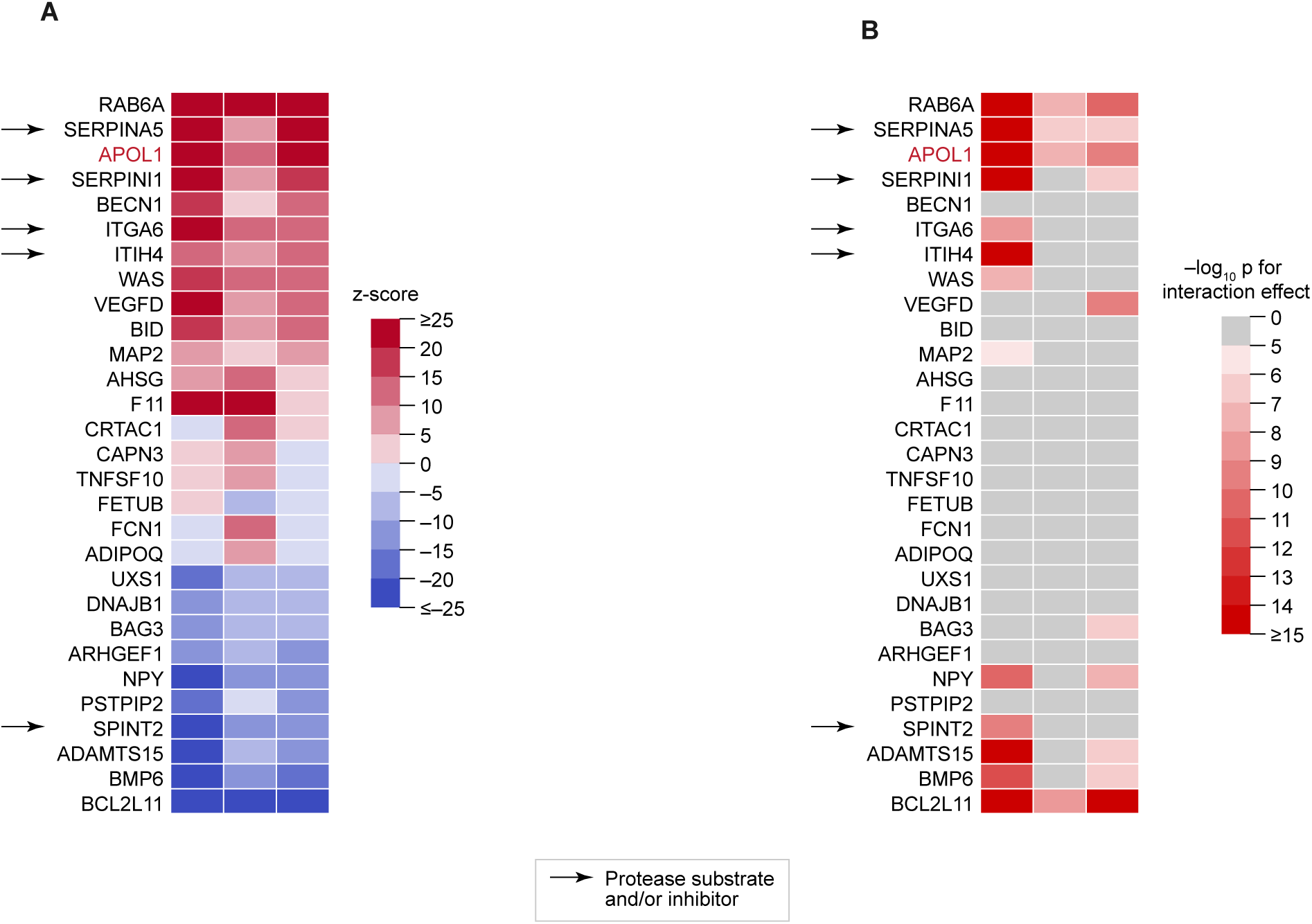
Variants in *KLKB1*, *F12* and *KNG1* are associated with plasma levels of multiple protease substrates and inhibitors. **(A)** Heatmap showing the associations between rs3733402 (*KLKB1*), rs1801020 (*F12*), and rs5030062 (*KNG1*) and the plasma levels of 29 proteins in individuals with European ancestry from the UK Biobank (Methods). Cell color indicates the signed z-score of the association test. Only proteins ranking in the top 20 strongest associations for at least one of the three variants are shown. **(B)** Heatmap showing the significance of the rs3733402 (*KLKB1*) x rs1801020 (*F12*), rs1801020 (*F12*) x rs5030062 (*KNG1*), and rs5030062 (*KNG1*) x rs3733402 (*KLKB1*) genetic interactions for plasma levels of the 29 proteins from (A). Cell color indicates the –log10 of the interaction p-value. Interactions with p > 10^−5^ are shown in grey. (A–B) The F12 and KLKB1 proteins themselves are omitted from the heatmaps for simplicity.

### *APOL1* and KKS variants do not show association with ESRD in the UK Biobank

Having established the platform-specific influence of *APOL1*, *KLKB1*, *F12*, and *KNG1* variants on Olink and SomaLogic measurements of plasma APOL1, we investigated whether these variants may also be relevant to APOL1-associated pathologies, specifically kidney disease. We used the algorithmically-defined outcome of end-stage renal disease (ESRD) (Herrington *et al*., 2017) as the most severe form of nephropathy reported in the UK Biobank with 921 and 44 cases identified among participants of European and African ancestry, respectively. Despite the established and consistently replicated increase in ESRD risk associated with the *APOL1* G1/G2 variants, we were unable to detect this association in UK Biobank participants with African ancestry (p > 0.05). Similarly, we did not observe an ESRD association for the *APOL1* V1 variant and the KKS variants in African or European participants (p > 0.05).

Interestingly, we observed that, regardless of APOL1 genotype, higher plasma APOL1 levels, as measured by both mass spectrometry (Health ABC) and Olink (UK Biobank), are associated with a reduced risk of all-cause mortality (**Fig. S8**). This association was independent of participants’ age and genetic ancestry (**Fig. S8**). Furthermore, in the UK Biobank, higher plasma APOL1 levels correlate with better kidney function and a lower risk of CKD (**Fig. S8**).

## Discussion

Early studies reported no correlation between circulating APOL1 levels and its G1/G2 mutations (Bruggeman *et al*., 2014; Weckerle *et al*., 2016; Andrews *et al*., 2022), which confer resistance to *T. b. rhodesiense* and significantly increase CKD risk (Friedman and Pollak, 2020). As a result, variation in plasma APOL1 levels received limited attention in parasitology and nephropathology studies. Recent advances in proteomic technologies applied to large plasma biobanks offer an opportunity to re-examine plasma APOL1 in the context of disease-associated mutations and other genetic factors. In this study, we analyzed plasma APOL1 measurements from four independent biobanks (UK Biobank, AASK, deCODE, and HealthABC) using three proteomic technologies (Olink, SomaLogic, and mass spectrometry). We show that APOL1 G1/G2 mutations act as causal *cis*-pQTLs for plasma APOL1 when measured by Olink and SomaLogic, but the effects estimated by the two technologies are directionally opposite (**Fig. 1**, **2A**). This platform dependency is mirrored, albeit with less strength, by the APOL1 V1 mutation, predicted to be structurally proximal to G1/G2 (**Fig. 4D**). Notably, none of these APOL1 variants (G1, G2, V1) impact plasma APOL1 levels measured by mass spectrometry (**Fig. 2B**, **4C**). Furthermore, we found that LOF variants in the kallikrein-kinin system (KKS) act as causal *trans*-pQTL for plasma APOL1 when measured by Olink but not SomaLogic (**Fig. 3A**).

Importantly, our findings on platform-dependent pQTL effects of the APOL1 V1 mutation and KKS variants are independently corroborated by recent data from the Multi-Ethnic Study of Atherosclerosis (MESA), which collected Olink and SomaLogic measurements for ∼2,000 plasma proteins (Nicholas *et al*., 2025). The MESA data confirm that the APOL1 V1 variant exhibits directionally opposite pQTL effects between APOL1_Olink_ and APOL1_SomaLogic_, with no effect on APOL1_MS_ (Nicholas *et al*., 2025). The MESA data also support *KLKB1* and *F12* variants acting as trans-pQTLs for APOL1_Olink_ but not APOL1_SomaLogic_ (Nicholas *et al*., 2025).

A plausible explanation for the platform-specific pQTL effects of APOL1 mutations is that these mutations induce protein conformational changes, which are differentially recognized by the reagents employed by Olink and SomaLogic – the so-called “epitope effect” (Ferkingstad *et al*., 2021; Eldjarn *et al*., 2023). Specifically, Olink antibodies may have higher affinity for the mutated APOL1 compared to the reference protein, whereas SomaLogic aptamers may have lower affinity. In individuals carrying the G1, G2, or V1 mutations, such differential affinity would result in apparently higher APOL1_Olink_ levels and apparently lower APOL1_SomaLogic_ levels, which would be consistent with our observations.

While we were unable to directly test the epitope hypothesis due to the proprietary nature of Olink reagents, several lines of indirect evidence support this possibility. The epitope hypothesis predicts that mass spectrometry, as a peptide-based rather than affinity-based technology, would be insensitive to protein conformational changes. This prediction aligns with our findings that G1, G2, and V1 mutations are not pQTLs for plasma APOL1_MS_ levels (**Fig. 2B**, **4C**). Consistent with our results, numerous published studies using enzyme-linked immunosorbent assay (ELISA), Western blotting, and mass spectrometry have also reported no association between G1/G2 mutations and plasma APOL1 levels (Bruggeman *et al*., 2014; Weckerle *et al*., 2016; Andrews *et al*., 2022). Biophysical analyses and molecular modeling have shown that G1/G2 variants alter the conformational dynamics and membrane interaction properties of the APOL1 C-terminal domain (Madhavan *et al*., 2017; Madhavan and Buck, 2021). The G1/G2 variants have also been reported to alter the avidity of an APOL1-targeting antibody (Weckerle *et al*., 2016). Finally, the APOL1 5’-UTR variant (V2) and the HPR variants (**Table 1**) show consistent, positively correlated pQTL effects across Olink and SomaLogic platforms (**Fig. 3B–C**), further supporting the hypothesis that platform discrepancies arise specifically from protein altering variants.

Proteolytic activity within the KKS cascade could also impact APOL1 structure or conformation, potentially explaining the platform-specific pQTL effects observed for KKS variants. If KKS proteases cleave APOL1 and Olink antibodies preferentially detect the cleaved protein form, then higher KKS activity, associated with functional alleles in *KLKB1*, *F12*, and *KNG1*, would yield apparently higher APOL1_Olink_ levels. Conversely, LOF alleles reducing KKS activity would result in apparently lower APOL1_Olink_ levels. This mechanism would have no impact on APOL1_SomaLogic_ measurements if SomaLogic aptamers detect both cleaved and uncleaved APOL1 forms equally. Under this hypothesis, the predicted associations of KKS variants with both APOL1_Olink_ and APOL1_SomaLogic_ levels would be consistent with our observations.

It is currently unknown if APOL1 is a direct substrate of plasma kallikrein or factor XIIa. However, we observed that LOF variants in *KLKB1*, *F12*, and *KNG1*, and their GxG interactions, are associated with altered plasma levels of known protease substrates and inhibitors, such as SERPINA5, SERPINI1, and ITIH4 (**Fig. 7**). These observations raise the possibility that APOL1 is similarly affected by KKS activity. Intriguingly, kallikrein and other hemostatic factors have been previously implicated in kidney disease (Yu *et al*., 1998, 2000; Tin *et al*., 2015), but the biological mechanisms underlying the connection remain unclear.

Despite prior evidence linking *APOL1* G1/G2 variants to kidney disease and their strong pQTL effects on plasma APOL1 observed in our study, we did not detect a significant association between these variants and ESRD risk in the UK Biobank. This null finding is likely attributable to low statistical power stemming from the low incidence of ESRD cases (0.65%) among participants of African descent in UK Biobank. According to our power analysis, this population has 90% power to detect G1/G2 ESRD odds ratios (ORs) greater than 2 (**Fig. S7**). For comparison, a study in the hospital-based Penn Medicine Biobank (PMBB), which reported a higher ESRD incidence of 8.4%, successfully detected a significant association between G1/G2 variants and ESRD with an OR of 1.6–1.7 (Bajaj *et al*., 2020; Zhang *et al*., 2024).

Compared to G1/G2, the *APOL1* V1 mutation and the KKS variants exhibit substantially weaker pQTL effects (e.g., β_G2_ = 1.2, posterior β_V1_ = 0.18 in UK Biobank AFR; **Fig. 1B**, **4B**). Working under the critical but unproven assumption that pQTL effect magnitude may translate proportionally to ESRD risk, we were not surprised by the absence of a significant association between V1/KKS variants and ESRD. Our power analysis estimates that the UK Biobank population of European descent has 90% power to detect V1 ESRD OR greater than 1.44 (**Fig. S7**). If we speculatively extrapolate ESRD ORs for *APOL1* V1 and KKS variants based on the assumption of proportionality to G1/G2, these projected ORs would be substantially lower than this estimated minimal detection threshold. Our inability to observe a V1 ESRD effect in UK Biobank is consistent with the PMBB study, which also failed to detect a significant V1 ESRD effect, independent of G1/G2 (Zhang *et al*., 2024). In contrast, prior case-control studies in African Americans with diabetic and non-diabetic nephropathies successfully reported a significant association between the V1 G allele and increased disease risk, independent of G1/G2 (Bostrom *et al*., 2012; Bailey *et al*., 2014). Taken together, these findings suggest that, although variant effects on plasma APOL1 are detectable in population-based cohorts, their clinical relevance may only emerge in disease-focused populations.

Interestingly, the observed pQTL effects of G1/G2 mutations on plasma APOL1 present several fascinating parallels to their reported impact on APOL1-associated pathologies. For instance, we show that the G1/G2 pQTL effects follow an additive model, where each copy of the risk allele contributes to an increase in the apparent protein level (**Fig. 1B–C**). This additive behavior aligns with the most recent study of CKD in West Africans, which, in contrast to prior studies suggesting recessive inheritance, reported an intermediate disease risk for individuals heterozygous for G1 or G2, compared to the homozygotes (Gbadegesin *et al*., 2025). The same study also reported that G2 carriers exhibit higher odds of disease compared to G1 carriers, consistent with the stronger pQTL effect of G2 relative to G1 in our analyses (**Fig. 1B–C**). We also show that, consistent with a prior report (Adamson *et al*., 2024), the G2 pQTL effect is attenuated by the p.N264K (M1) variant (**Fig. S2F**), a finding that correlates with a protective effect of M1 on G2-associated risk of kidney disease (Gupta *et al*., 2023) and *Trypanosoma* infections (Cuypers *et al*., 2016; Cooper *et al*., 2017). These similarities between G1/G2 pQTL effects and disease risk hint at the possibility that affinity-based proteomic assays may capture functionally relevant aspects of APOL1 biology, even when changes in total protein abundance are not involved.

Experimental evidence is needed to validate the hypothesis that G1/G2/V1 mutations alter APOL1 structure and to understand how such changes might compare to those potentially driven by KKS activity. It is also important to consider that plasma APOL1 does not circulate freely but is transported by specific HDL particles, namely trypanosoma lytic factors 1 (TLF-1) and 2 (TLF-2). Therefore, any structural changes in APOL1, arising from either mutations or KKS, may also impact the protein’s interaction with its HDL carriers. Given the strong pathogenicity of G1/G2 variants, defining their impact on APOL1 structure and identifying other factors with potential structural effects, such as the KKS pathway, is crucial for understanding disease etiology and generating new therapeutic hypotheses.

## Methods

### Study populations

The UK Biobank (UKB) is a prospective cohort of over 500,000 participants aged 40–69 at recruitment with extensive genotype and phenotype data, as detailed at https://biobank.ndph.ox.ac.uk/showcase/. In the Pharma Proteomics Project (PPP), plasma samples from ∼50,000 participants were analyzed using the Olink Explore 3072 platform to measure relative abundance of ∼3,000 circulating proteins (Sun *et al*., 2023). The UKB-PPP cohort was used for pQTL association tests, whereas the entire UKB cohort was used for disease association tests. All participants provided informed consent.

We used the Pan-UKB consortium dataset (Karczewski *et al*., 2024) to assign genetic ancestry to individuals in the UKB-PPP cohort, focusing on those of European and African descent. To minimize relatedness within each sample set, we randomly excluded one individual from each pair with a kinship coefficient (Pi_hat) greater than 0.125. This relatedness filter, while less stringent than standard approaches, maximized sample size, yielding to 47,178 and 948 individuals of European and African ancestry, respectively. Nearly 100% of these individuals (47,090 and 947) had both genotype data and APOL1 measurements, whereas 45,444 (96%) and 893 (94%) had both exome sequences and APOL1 measurements.

The Health, Aging, and Body Composition (Health ABC) study is an observational cohort study investigating the functional decline of healthy aging individuals (Newman *et al*., 2023). The cohort includes 3,075 participants aged 70–79, with 33% men and 45% women self-reported as African-American. Mass spectrometry and the Olink assay were performed on a random subset of 1,094 and 560 Health ABC participants, respectively **(Table S1**).

### UKB: Genotype data processing and quality control

Genotyping, imputation and quality control of UKB samples were performed in the consortium pipeline (Bycroft *et al*., 2018). We used version 3 of the imputed data, which was generated using the Haplotype Reference Consortium (HRC) and the UK10K haplotype resource as the imputation panel (described in the UK Biobank Category 100319). In addition to the baseline QC practices implemented by UKB, variants with an INFO score below 0.6 were filtered out. Exome data were also processed through the UKB consortium pipeline and accessed via the UK Biobank Research Analysis Platform (UKB-RAP). Details of exome sequencing procedures are available on the UKB website (https://dnanexus.gitbook.io/uk-biobank-rap/science-corner/whole-exome-sequencing-oqfe-protocol) as well as related publications (Backman *et al*., 2021; Okada and Wang, 2021). Within each sample set (defined above), variants with Minor Allele Count (MAC) below 10 were filtered out.

Genotypes from both imputed and genotyped data (hg19) were lifted over to hg38 using the Genome Analysis Toolkit (GATK) LiftoverVcf tool (https://gatk.broadinstitute.org/hc/en-us). Variants without unique mapping were filtered out. Exome data were already provided in the hg38 form.

Association tests were performed using primarily imputed genotype data, with the exception of the APOL1 G1 variant in the African population. Comparison of exome and imputed genotype data revealed discrepancies in G1M and G1G variant calls. Specifically, over 20% (73 of 351) of heterozygous G1M carriers and 5% (17 of 349) of heterozygous G1G carriers identified by exome sequencing were misidentified as homozygous in the imputed genotype data (**Fig. S1A-B**). Further supporting imputation inaccuracies, we observed a low correlation between G1G and G1M genotypes (60 samples showed discordant genotypes for G1G and G1M; **Fig. S1C**), excess linkage disequilibrium with G2 (56 samples appeared to be homozygous for G1M and heterozygous for G2, an unlikely genotype pattern given that G1 and G2 never occur on the same haplotype; **Fig. S1D**) and deviation from Hardy-Weinberg equilibrium in the imputed data (p < 10^−14^) compared to the exome data (p > 0.05; **Fig. S1E**). As a result of these discrepancies, we prioritized exome data when available. We confirmed that the association between G1/G2 and plasma APOL1 levels was similarly likely to be causal in the exome data alone, imputed genotype data alone, or a combination of both (**Fig. S1F-G; Fig. 1A**).

For visualization of individual variant effects (e.g., in violin plots or kernel density estimation plots), exome data were prioritized, unless the variant was absent (e.g., the non-coding HPR lead cis-variant).

### UKB-PPP: pQTL analyses

In the UKB-PPP dataset, plasma protein levels are reported as Olink’s Normalized Protein eXpression (NPX) values, which reflect relative change in protein abundance on a log_2_ scale following plate and batch normalization (Sun *et al*., 2023).

Association tests between genotypes and protein levels were performed using tensorQTL v1.0.9 (https://github.com/broadinstitute/tensorqtl). Prior to analysis, protein NPX matrices were inverse-normal transformed within each study population. Age, sex, five genotype PCs and ten protein expression PCs were included as covariates. Genotype PCs were provided by the Pan-UKB consortium. Protein expression PCs were calculated using the PCA function of scikit-learn applied to the inverse-normal transformed protein NPX matrix.

Genetic effects were estimated using an additive model, where each copy of the effect allele contributes equally to the change in protein levels. To facilitate interpretation, variant effect sizes (β) are reported as per-allele changes in raw (rather than inverse-normal transformed) protein NPX values, unless explicitly stated. For variants in linkage disequilibrium with the APOL1 G2 variant, which exhibits a large effect on plasma APOL1 levels in the Olink-based measurements, we report posterior β estimates from the FINEMAP algorithm (Benner *et al*., 2016). This approach accounts for the strong effect of G2 and provides a more accurate estimate of the independent effects of other variants in the region (**Fig. S2A–E**). Marginal and posterior β are explicitly distinguished throughout the manuscript. Effect sizes from the replication cohorts (AASK and deCODE; SomaLogic-based measurements) are reported as marginal β, since posterior β estimates are not available. Given that the effect of G2 in these studies is much smaller than in UKB, we do not anticipate marginal β to introduce substantial bias.

### UKB: fine-mapping

Fine-mapping of each locus associated with protein expression was performed using FINEMAP v.1.3.1 (Benner *et al*., 2016) and susieR v.0.11.43 (Wang *et al*., 2020) with default parameter settings. For genes on the plus strand, the *cis*-locus was defined as the region spanning 0.2 Mb upstream of the transcription start site (TSS) to 0.2 Mb downstream of the transcription end site (TES). For genes on the minus strand, the *cis*-locus was defined as the region spanning 0.2 Mb upstream of the TES to 0.2 Mb downstream of the TSS. Input data included summary statistics from tensorQTL and the in-sample LD matrix adjusted for the same covariates used in the association tests. A maximum of ten causal variants was allowed.

### UKB: *HPR* copy number analysis

HPR copy number (CN) data was obtained from the UKB-RAP, which provides CN estimations derived from whole-exome sequencing data analyzed by the Illumina DRAGEN platform (https://developer.illumina.com/dragen). For association tests and fine-mapping of the *HPR* region, HPR CN values were included as an additional variable, mean-centered and scaled like other genetic variants.

### UKB: testing for genetic interactions

To identify potential genetic interactions between pairs of variants, we employed a linear regression model using the statsmodel package in Python. For each pair of variants (*v*_1_ and *v*_2_), we fit the following model: *y* ∼ *v*_1_ + *v*_2_ + *v*_1_ × *v*_2_ + *covariates*, where *y* represents the phenotype, and *covariates* represents the same set of variables used in the association tests (see above). The significance of the genetic interaction was assessed using the p-value of the interaction term.

To assess the difference in the effect of variant *v*_1_ conditioned on the presence of another variant (*v*_2_), we employed a stratified analysis using separate linear regression models. Two models (M1 and M2) were fit: *y* ∼ *v*_1_ *covariates* when *v*_2_ is absent (M1), and *y* ∼ *v*_1_ + *covariates* when *v*_2_ is present (M2). Let β_1_ and β_2_ denote the estimated effects of *v*_1_ in models M1 and M2, respectively, with corresponding standard errors σ and σ. We calculated the t-statistic for the difference in effects as 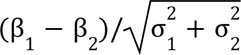. Under the null hypothesis of no difference in effects, this statistic is assumed to follow a t-distribution with *N*_1_ + *N*_2_ – 4 degrees of freedom, where *N*_1_ and *N*_2_ are the sample sizes in the respective strata.

When conditioning on the p.N264K variant, both the heterozygous (REF/ALT) and the homozygous alternative (ALT/ALT) genotypes for *v*_2_ were included in model M2. When conditioning on the G1 or G2 variants, only the REF/ALT genotype was included in M2. This decision was based on the observation that G1 and G2 are never found on the same haplotype, such that, if one variant in ALT/ALT, the other is always REF/REF. As a result, including the ALT/ALT genotype in M2 would only reduce power.

### Health ABC: proteomic analyses

Mass spectrometry analysis was performed on a random set of 1,111 Health ABC participants using samples collected at baseline and second year follow up visits. Data from the two visits were averaged to produce a single value per individual. Samples that failed during processing or data quality control were excluded from further examination, resulting in 1,094 participants available for downstream analysis.

#### Materials

HEPES and EPPS buffers were obtained from Boston BioProducts (Milford, MA, USA), and cOmplete ULTRA and PhosStop inhibitor tablets were obtained from Roche (Mannheim, Germany). Trypsin/Lys-C Mix was obtained from Promega Corporation (Madison, WI, USA). Tandem mass tag (TMT) 16-plex reagent sets were obtained from Thermo Fisher Scientific (Rockford, IL). Unless otherwise stated, all other chemicals were purchased from Sigma-Aldrich (St. Louis, MO, USA).

#### Sample preparation for liquid chromatography-mass spectrometry (LC-MS)

The samples were prepped using our in-house automated multiplexed proteome profiling platform (AutoMP3) protocol (Gaun *et al*., 2021). Neat plasma (2 µL) was diluted with 38 µL of lysis buffer (75 mM NaCl, 50 mM HEPES pH 8.5, 3% sodium dodecyl sulfate (SDS) with cOmplete ULTRA and PhosSTOP tablets). The samples were reduced with 5 mM final concentration dithiothreitol (DTT) for 30 minutes at 25°C, then alkylated with 15 mM final concentration iodoacetamide in the dark for 30 minutes at 25°C. Excess iodoacetamide was quenched with 5 mM final concentration DTT and incubated in the dark for 15 minutes at 25°C. Samples were cleaned up using a mixture of Sera-Mag beads (Cytiva Life Sciences, 44152105050350, 24152105050350; Marlborough, MA, USA). Beads were combined 1:1 (E3:E7) then washed 3 times with water. Samples were added directly to the beads, then acetonitrile was added to a final concentration of 75% and the mixture was incubated for 20 minutes at room temperature for protein binding. Beads were immobilized on a magnet then washed two times with 70% ethanol (200 µL) followed by two washes with 100% acetonitrile (ACN, 200 µL). The beads were resuspended in 40 µL of digestion buffer (50 mM EPPS pH 8.5, 10 mM CaCl2) then 10 µL of Trypsin/Lys-C (0.33 mg/mL) was added. The proteins were digested on-bead for 1 hour at 37°C, 1000 rpm. Digested samples were immobilized on a magnet and the peptide supernatant was transferred to a new plate. To enable analysis across multiple plexes, a bridge sample was created by pooling a small amount of each peptide sample. Tandem mass tag (TMT) labeling was performed using 30 µg of peptides and adding 0.5 mg of TMT reagent. The TMT reaction was incubated for 1 hour at 25°C then excessed TMT was quenched with 0.3% final concentration hydroxylamine. For each plex, samples were combined, mixed, then split into several wells (≤100 µL per well) for peptide cleanup. For peptide binding, acetonitrile was added to a final concentration of 95%. Beads were immobilized on a magnet then washed one time with 100% ACN (1 mL). Peptides were eluted using 3 rounds of 5% ACN (100 µL), then plexes were re-combined and dried down in a SpeedVac overnight at 25°C. The combined samples were resuspended in 5% ACN / 5% formic acid (FA) then analyzed on a mass spectrometer.

#### Liquid chromatography-mass spectrometry

Peptides were analyzed on an Orbitrap Eclipse mass spectrometer coupled to an Ultimate 3000 (Thermo Fisher Scientific). Peptides were separated on an IonOpticks Aurora microcapillary column (75 μm inner diameter, 25 cm long, C18 resin, 1.6 μm, 120 Å, Gen2). The total LC-MS run length for each sample was 185 min including a 165 min gradient from 6 to 35% ACN in 0.1% formic acid. The flow rate was 300 nL / min, and the column was heated at 60° C. A Field Asymmetric Ion Mobility Spectroscopy device (FAIMS Pro) was connected to the Orbitrap Eclipse mass spectrometer and a method with four experiments, where each experiment utilized a different FAIMS Pro compensation voltage: −40, −50, −60, and −70 volts was used. Each of the four experiments had a 1.25 second cycle time. A high resolution MS1 scan in the Orbitrap (m/z range 400-1,600, 120k resolution, standard Automatic Gain Control (AGC) target settings, “Auto” max injection time, ion funnel RF of 30%) was collected from which precursors were selected for MS2 followed by SPS MS3 analysis. For MS2 spectra, ions were isolated with the quadrupole mass filter using a 0.7 m/z isolation window. The MS2 product ion population was analyzed in the quadrupole ion trap (CID, “standard” AGC target, normalized collision energy 35%, “Custom” max injection time set to 35 ms) and the MS3 scan was analyzed in the Orbitrap (HCD, 50k resolution, normalized collision energy 45%, “custom” AGC target with normalized AGC target set to 200% and max injection time 200 ms). Up to ten fragment ions from each MS2 spectrum were selected for MS3 analysis using SPS. A human database was used for Real Time Search (RTS) (Schweppe *et al*., 2020) and a maximum of two missed cleavages and one variable modification was allowed and FDR filtering enabled. The maximum search time was set to 35 ms and a Xcorr of 1, dCN 0.1 and precursor of 10 ppm for charge state 2, 3 and 4 was used.

#### Peptide Identification and Quantification

Mass spectrometry data were processed using a previously described software pipeline (Huttlin *et al*., 2010). Raw files were converted to mzXML files and searched against a human database (downloaded on October 2nd, 2020), in forward and reverse orientations using the Sequest algorithm (Eng, McCormack and Yates, 1994; Huttlin *et al*., 2010). Database searching matched MS/MS spectra with fully tryptic peptides from this composite dataset with a precursor ion tolerance of 20 ppm, and 0.6 Da product ion tolerance. Static carbamidomethylation modifications of cysteine residues (+57.02 Da) and TMTpro tags on peptide N-termini and lysines (+304.20 Da) were used. Oxidation of methionine (+15.99 Da) was set as a variable modification. Linear discriminant analysis was used to filter peptide spectral matches to a 1% FDR (false discovery rate) as described previously (Elias and Gygi, 2007). Non-unique peptides that matched to multiple proteins were assigned to proteins that contained the largest number of matched redundant peptide sequences using the principle of Occam’s razor. Quantification of TMT reporter ion intensities was performed by extracting the most intense ion within a 0.003 m/z window at the predicted m/z value for each reporter ion. Relative abundances of peptides were calculated as log2 ratio of sample to bridge, where bridge sample was a common control shared across all TMT plexes.

#### Olink Platform

Olink analysis was performed on 658 samples selected at random from those chosen for mass spectrometry. Samples were shipped to Olink Proteomics (Waltham, MA, USA) and processed by the manufacturer. Normalized protein expression (NPX) values were generated after quality control, normalization, and batch effect correction. Samples that failed during processing or data quality control were excluded from further examination, resulting in 560 samples available for downstream analysis.

### Health ABC: inference of G1, G2 and V1 genotypes from peptide sequence

To infer APOL1 variant status, we analyzed peptide-level mass spectrometry data (**Fig. S3**) (Norris Bradley *et al*., 2020).

For rs2239785 (V1), we utilized two peptides, SELEDNIRR and LEDNIRR, which overlap the V1 variant position and correspond to the REF (G) and ALT (A) alleles, respectively. A scatter plot of the two peptide measurements was used to infer genotypes (**Fig. S3B**). Samples with two copies of the reference allele (G/G) were expected to fall along the y=0 line, those with two copies of the alternative allele (A/A) along the x=0 line, and heterozygotes (G/A) along the y=x line. We classified samples as G/G, A/A or G/A, depending on whether they fall above, below or within a defined range (x–0.5 < y < x+1; **Fig. S3B**). Samples with a peptide measurement of exactly 0 were classified as homozygous for the corresponding allele, regardless of whether they fall within the defined range (**Fig. S3B**).

For G1/G2, we utilized the peptide LNILNNNYK, which overlaps the G1M/G2 variants and is theoretically only observed in individuals with a REF/REF or a REF/ALT genotype for both. This peptide’s abundance is expected to be comparable to other APOL1 peptides in REF/REF individuals, but substantially lower in REF/ALT and ALT/ALT individuals. We trained a lasso model (scikit-learn v1.3.0, alpha=0.01 and default settings for all other parameters) in European samples (virtually all REF/REF) to predict this peptide’s abundance using other 18 APOL1 peptides as features. The model was then applied to African samples to compare predicted and observed peptide measurements. We classified African samples as REF/ALT or ALT/ALT, depending on whether their observed values fell below y < 5/3*x or y < 5/3*x-1 thresholds, respectively (where x is the predicted value; **Fig. S3A**). Samples with a peptide measurement of exactly 0 were additionally classified as ALT/ALT (**Fig. S3A**).

### Comparison to AASK and deCODE data

Plasma pQTL summary statistics for the deCODE study were obtained from https://www.decode.com/summarydata/. Plasma pQTL summary statistics for the AASK study were obtained from (Surapaneni *et al*., 2022). Throughout the manuscript, we report the effect sizes of the alternative alleles as defined in hg38, regardless of the allele frequencies, unless otherwise noted. For example, in the case of V1, we report the effect size of the reference allele (G) as, similar to G1/G2 mutations, it is substantially more common in populations of African ancestry than in Europeans.

### UKB: disease associations

The UKB field ID 42026, which reports the date of end-stage renal disease (ESRD) diagnosis, was used as evidence of disease. The field is based on a combination of multiple factors including health records, self-reports, and death certificates. We binarized the field, assigning a value of 1 to individuals with an ESRD diagnosis and 0 to those without a diagnosis (or with missing data). A relatedness filter, as defined by the Pan-UKB consortium, was applied. We then performed logistic regression using disease diagnosis as an outcome and including age, sex and 10 genotype PCs as covariates.

### UKB: adjusting for blood pressure-related factors

To assess the impact of *KLKB1*, *F12* and *KNG1* variants on APOL1 levels while controlling for blood pressure-related factors, we performed linear regressions which included age, sex, five genotype PCs and the following covariates: diastolic blood pressure (UKB field 4079), calculated as the mean of two repeated measurements; systolic blood pressure (UKB field 4080), calculated as the mean of two repeated measurements; history of hypertension diagnosis (UKB field 2966), coded as a binary variable (1 = diagnosis at any age, 0 = no diagnosis); history of blood pressure medication use (UKB fields 6153 and 6177), coded as a binary variable (1 = medication record present, 0 = no medication record). Missing values for any covariate were imputed using the sample mean. Linear regression models were fit without regularization.

### Power calculation

To evaluate the statistical power to identify associations between risk alleles and ESRD phenotype in our UKB dataset, we conducted a power analysis using a two-proportion z-test (Sham and Purcell, 2014). The test compares the proportion of individuals with the high-risk genotype (under recessive model) in cases (individuals with ESRD) and controls (individuals without ESRD).

Given the number of cases and control (which we directly obtain in each population group as defined in the Pan-UKB), we calculated the probability of detecting a statistically significant difference in high-risk genotype frequencies between these groups with the type 1 error rate of 0.05. Within a population, the risk allele frequencies were calculated as (2 x number of individual with risk homozygous genotype + number of individuals with heterozygous genotype) / (2 x number of total individuals).

### Statistical analysis

All the statistical tests were two-sided. Reported p-values are unadjusted unless otherwise noted. Error bars represent standard errors of the mean (SEM) unless otherwise noted.

## Data and code availability

The UK Biobank (UKB) data were accessed following the processes described in https://www.ukbiobank.ac.uk/enable-your-research. The UKB Research Analysis Platform (RAP) was primarily used for UKB data analysis. The Health ABC data were accessed following the processes described in https://www.nia.nih.gov/healthabc-study. Plasma pQTL summary statistics for the deCODE study were obtained from https://www.decode.com/summarydata/. Plasma pQTL summary statistics for the AASK study were obtained from (Surapaneni *et al*., 2022). Variant-level summary statistics generated in this manuscript are available throughout the text and in the Supplementary Data 1 file available at (https://storage.googleapis.com/apol1-paper/APOL1_supplementary_data_1.tar.gz).

Software tools used for data analysis and visualization include: dx-toolkit (https://github.com/dnanexus/dx-toolkit); FINEMAP v.1.3.1 (http://www.christianbenner.com/); GATK v.4.1.9.0 LiftoverVcf (https://gatk.broadinstitute.org/); matplotlib v.3.3.4 (https://matplotlib.org); NumPy v.1.20.1 (https://numpy.org); pandas v.1.1.4 (https://pandas.pydata.org); scikit-learn v.0.24.1 (https://scikit-learn.github.io/stable); SciPy v.1.6.2 (https://scipy.org/); seaborn v.0.11.1 (https://seaborn.pydata.org); susieR v.0.11.43 (https://github.com/stephenslab/susieR); tensorQTL v.1.0.9 (https://github.com/broadinstitute/tensorqtl). The code used in this manuscript is available at https://github.com/QingboWang/apol1_manuscript.

## Acknowledgements

We would like to thank Anil Raj, Martin Mullis, Qi Hao, Manoj Rathinaswamy, Bernd Wranik, Melissa Pilling, Robert Cohen and Nick van Bruggen for useful discussions and feedback. We also acknowledge Steven Cummings and the Health ABC study for enabling validation analyses. Research with the Health ABC data was supported by National Institute on Aging (NIA) Contracts N01-AG-6-2101; N01-AG-6-2103; N01-AG-6-2106; NIA grant R01-AG028050, and NINR grant R01-NR012459. This research was funded in part by the Intramural Research Program of the NIH, National Institute on Aging. Research with the UK Biobank Resource was conducted under application number 18448. This study was funded by Calico Life Sciences LLC.

