## Supplementary Material for "Platform-dependent effects of genetic variants on plasma APOL1"

**Table S1. Demographic characteristics of the Health ABC sub-cohorts that were used for mass spectrometry and Olink analyses.**

| | Sub-cohort examined by mass spectrometry | Sub-cohort examined by Olink |
|---|---|---|
| Number of participants: n | 1094 | 560 |
| Age at baseline: mean (SD) | 73.96 (3.21) | 73.82 (3.14) |
| BMI: mean (SD) | 27.40 (4.55) | 27.72 (4.50) |
| Systolic blood pressure: mean (SD) | 136.23 (20.46) | 135.78 (20.04) |
| Cystatin C: mean (SD) | 1.03 (0.30) | 1.03 (0.29) |
| Vitamin D: mean (SD) | 25.97 (11.44) | 25.78 (10.85) |
| C-reactive protein (CRP): mean (SD) | 6.93 (27.06) | 8.36 (32.65) |
| Albumin/creatinine ratio in urine: mean (SD) | 41.38 (168.60) | 41.19 (187.01) |
| Creatinine: mean (SD) | 1.05 (0.32) | 1.04 (0.32) |
| Fasting glucose: mean (SD) | 102.39 (29.66) | 104.23 (31.02) |
| Hemoglobin A1C: mean (SD) | 6.26 (0.96) | 6.31 (1.02) |
| LDL: mean (SD) | 121.62 (33.39) | 122.40 (33.20) |
| Triglycerides: mean (SD) | 140.25 (89.96) | 144.02 (82.73) |
| HDL: mean (SD) | 52.58 (15.82) | 52.35 (16.46) |
| Gender (female): n (%) | 554 (50.6) | 286 (52.3) |
| Race (African): n (%) | 390 (35.6) | 194 (35.5) |
| Education level: n (%) | | |
| Less than high school | 262 (23.9) | 144 (26.3) |
| High school graduate | 344 (31.4) | 176 (32.2) |
| Postsecondary | 485 (44.3) | 226 (41.3) |
| Former smoker: n (%) | 550 (50.3) | 268 (49.0) |
| Chronic kidney disease (CKD): n (%) | 18 (1.6) | 10 (1.8) |
| Diabetes: n (%) | 147 (13.4) | 79 (14.4) |

**Table S2. Allele frequencies of kallikrein-kinin system (KSS) variants in the UK Biobank and the deCODE cohorts.**

| Gene | Variant | MAF (UK Biobank) | MAF (deCODE) |
|---|---|---|---|
| *KLKB1* | rs3733402 | 0.48848 | 0.49078 |
| *F12* | rs1801020 | 0.254797 | 0.25319 |
| *KNG1* | rs5030062 | 0.37349 | 0.35548 |

Figure S1

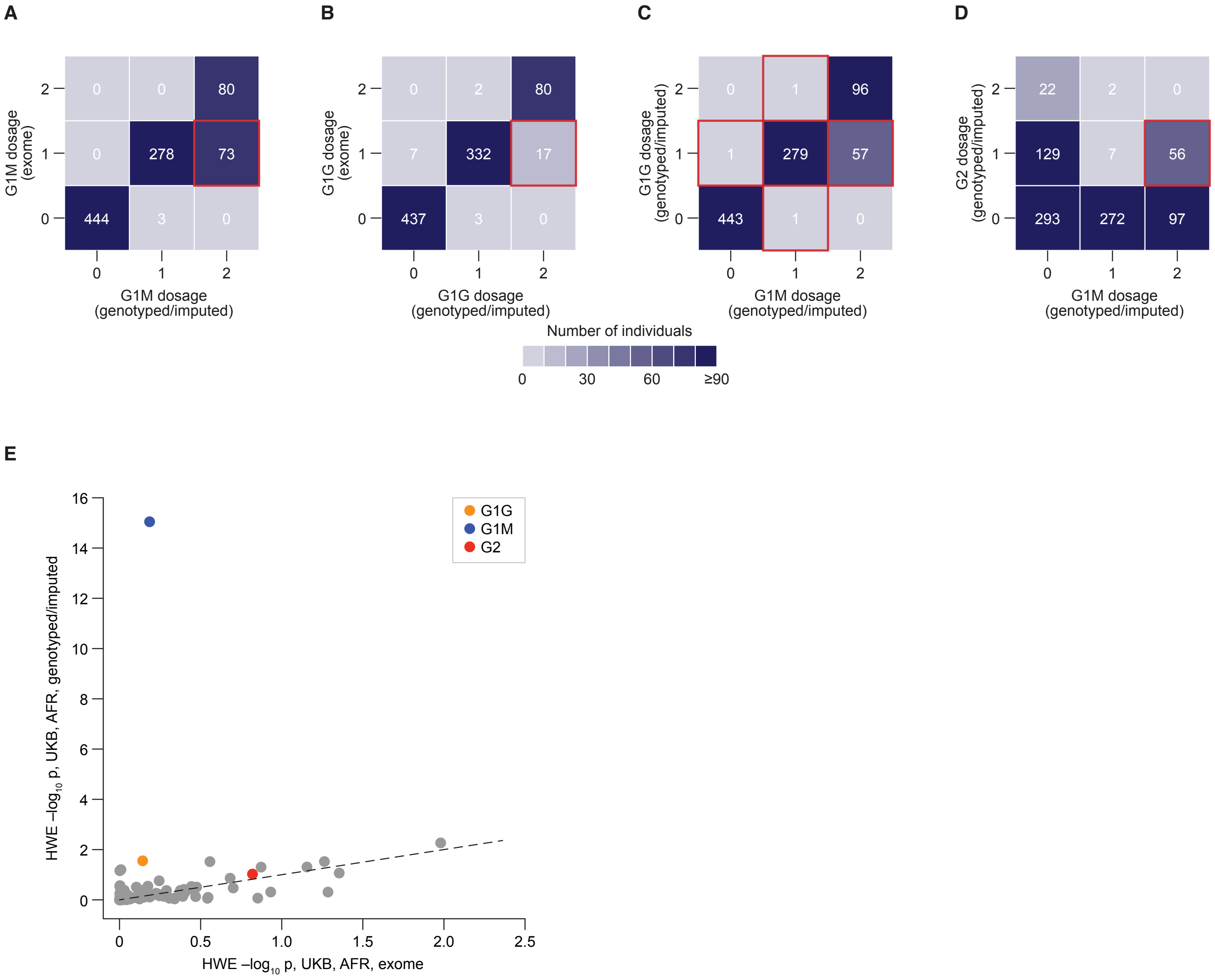


**Supplementary Figure 1. Exome sequencing reveals discrepancies in *APOL1* G1M imputation in the UK biobank.** Genotypes for the *APOL1* G1M, G1G and G2 variants were obtained for 878 UK Biobank-PPP participants with African ancestry (UKB, AFR) using both imputed genotype array data and exome sequencing. **(A–B)** Heatmaps comparing genotype calls for the G1M **(A)** and G1G **(B)** variants obtained from imputed (x-axis) and exome-derived data (y-axis). Each cell represents the number of individuals with the corresponding genotypes. Non-zero values off the diagonal represent conflicting genotype calls. For example, over 20% (73 of 351; **A**, red square) of heterozygous G1M carriers and 5% (17 of 349; **B**, red square) of heterozygous G1G carriers identified by exome sequencing are misidentified as homozygous in the imputed genotype data. **(C–D)** Heatmaps comparing genotype calls for the G1M (x-axis) and G1G (**C**; y-axis) or G2 (**D**; y-axis) variants obtained from imputed data. Each cell represents the number of individuals with the corresponding genotypes. Improbable genotype combinations are highlighted in red. For example, 60 individuals show discordant genotypes for G1G and G1M **(C)**. Similarly, 56 individuals appear to be homozygous for G1M and heterozygous for G2, an unlikely genotype pattern given that G1 and G2 never occur on the same haplotype **(D)**. **(E)** Scatter plot comparing deviation from Hardy-Weinberg Equilibrium (HWE) for *cis-APOL1* variants estimated from exome-derived data (x-axis) and imputed genotypes (y-axis). Each dot corresponds to a variant, with G1M being a clear outlier.

**Figure S2**

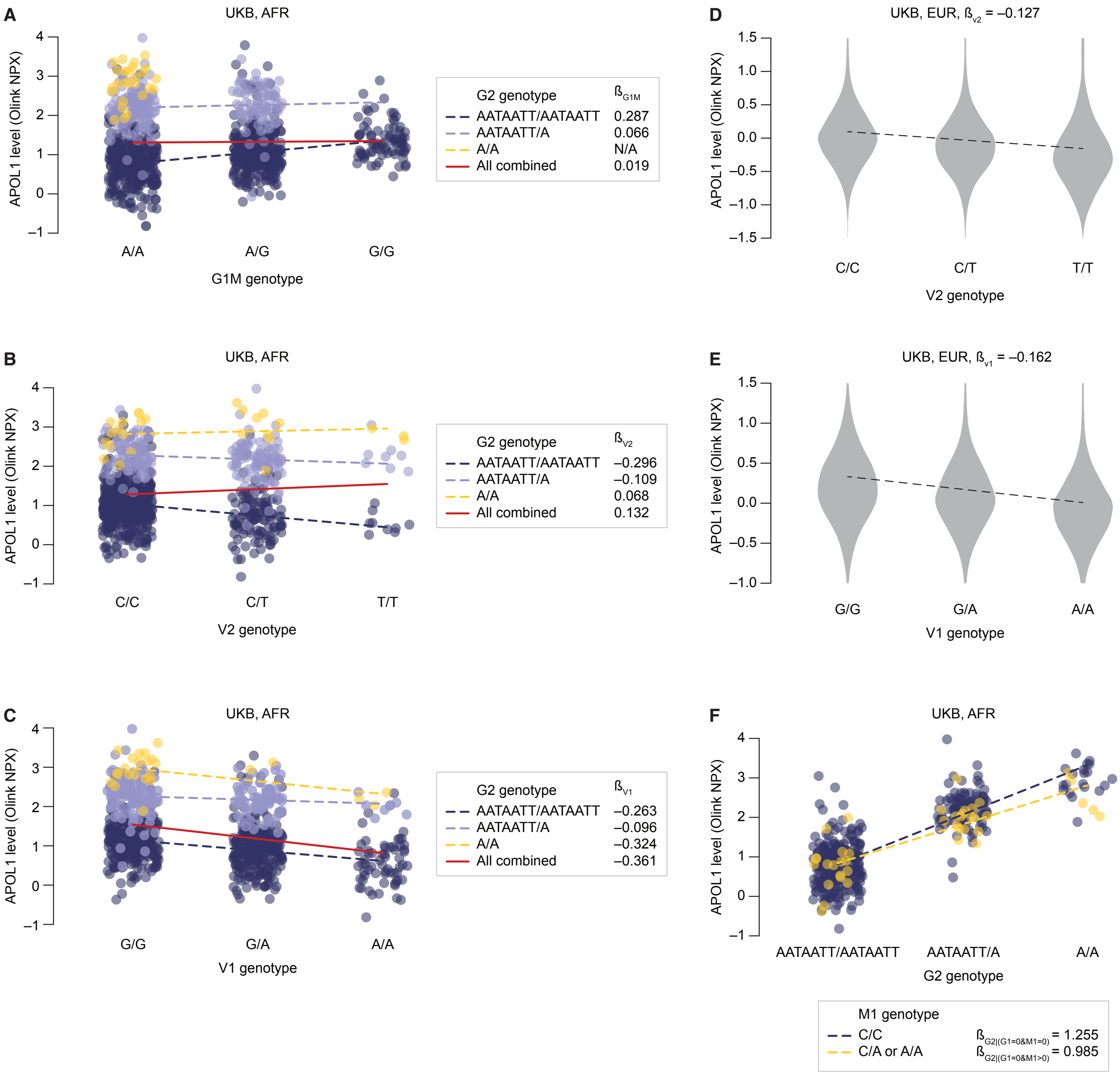


**Supplementary Figure 2. Linkage disequilibrium with G2 and genetic interactions with M1 influence the estimation of effect sizes for *APOL1* cis-variants. (A–C)** Scatter plots showing the effects of G1M **(A)**, V2 **(B)** or V1 **(C)** genotypes (x-axis) on plasma APOL1 levels (measured by Olink; y-axis) in UK Biobank participants with African ancestry (UKB, AFR), stratified by G2 genotype. Each data point represents an individual. Data points are colored according to G2 genotype: navy (REF/REF), light blue (REF/ALT) and yellow (ALT/ALT). Within each G1M **(A)**, V2 **(B)** or V1 **(C)** genotype group, data points are jittered along the x-axis for visualization purposes. Lines represent linear regression fits for each G2 genotype group (dashed, color-coded) and the overall trend (solid, red). **(D–E)** Violin plots showing the effects of V2 **(D)** or V1 **(E)** genotypes (x-axis) on plasma APOL1 levels (measured by Olink; y-axis) in UK Biobank participants with European ancestry (UKB, EUR). Dashed lines represent linear regression fits. **(F)** Scatter plot showing the effect of the G2 genotype (x-axis) on plasma APOL1 levels (measured by Olink; y-axis) in UK Biobank participants with African ancestry (UKB, AFR), stratified by M1 genotype. Each data point represents an individual. Data points are colored according to M1 genotype: navy (REF/REF), and yellow (REF/ALT or ALT/ALT). Within each G2 genotype group, data points are jittered along the x-axis for visualization purposes. Lines represent linear regression fits for each M1 genotype group (dashed, color-coded).

**Figure S3**

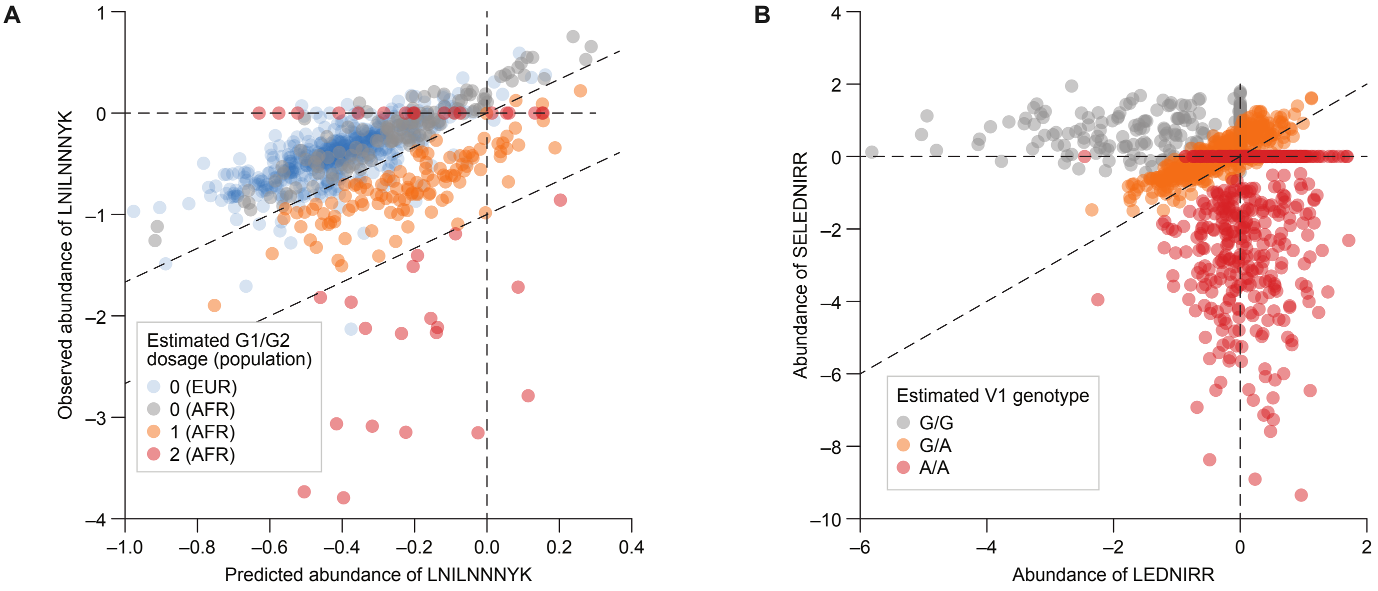


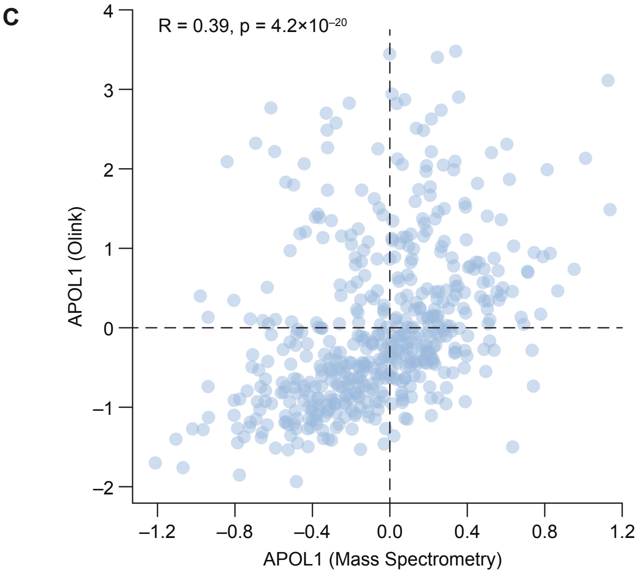


**Supplementary Figure 3. In Health ABC, *APOL1* G1/G2 and V1 variant genotypes can be inferred from peptide sequence analysis. (A)** Inference of G1/G2 ALT dosage from peptide sequence analysis in mass spectrometry (Methods). Each data point represents an individual. The dosage of the ALT alleles for G1M/G2 in each individual was estimated by comparing the predicted (x-axis) and observed (y-axis) abundance of the peptide LNILNNNYK, which overlaps both the G1M and G2 variants. Predictions were generated using a lasso model trained on G1/G2-independent peptides in individuals with European ancestry (virtually all REF/REF) and applied to individuals with African ancestry (Methods). Data points are colored according to their ancestry (blue for EUR, other colors for AFR) and inferred G1/G2 genotype (grey for REF/REF, orange for REF/ALT and red for ALT/ALT). Dashed lines indicate classification boundaries. **(B)** Inference of V1 ALT dosage from peptide sequence analysis in mass spectrometry (Methods). Each data point represents an individual. The dosage of the ALT allele for V1 in each individual was estimated by comparing the observed abundance of the LEDNIRR (x-axis) and SELEDNIRR (y-axis) peptides, which correspond to the V1 A (ALT) and G (REF) alleles, respectively. Data points are colored according to their inferred genotypes (grey for REF/REF, orange for REF/ALT and red for ALT/ALT). Dashed lines indicate classification boundaries. **(C)** Scatter plot comparing plasma APOL1 levels measured by mass spectrometry (x-axis) and Olink's platform (y-axis) in the same individuals from Health ABC. **(D)** Effect sizes (β) of the inferred G1/G2 and V1 genotypes on individual peptide level measurements in Health ABC. Age, sex, body mass index (BMI) and smoking status (ever/never) were included as covariates.

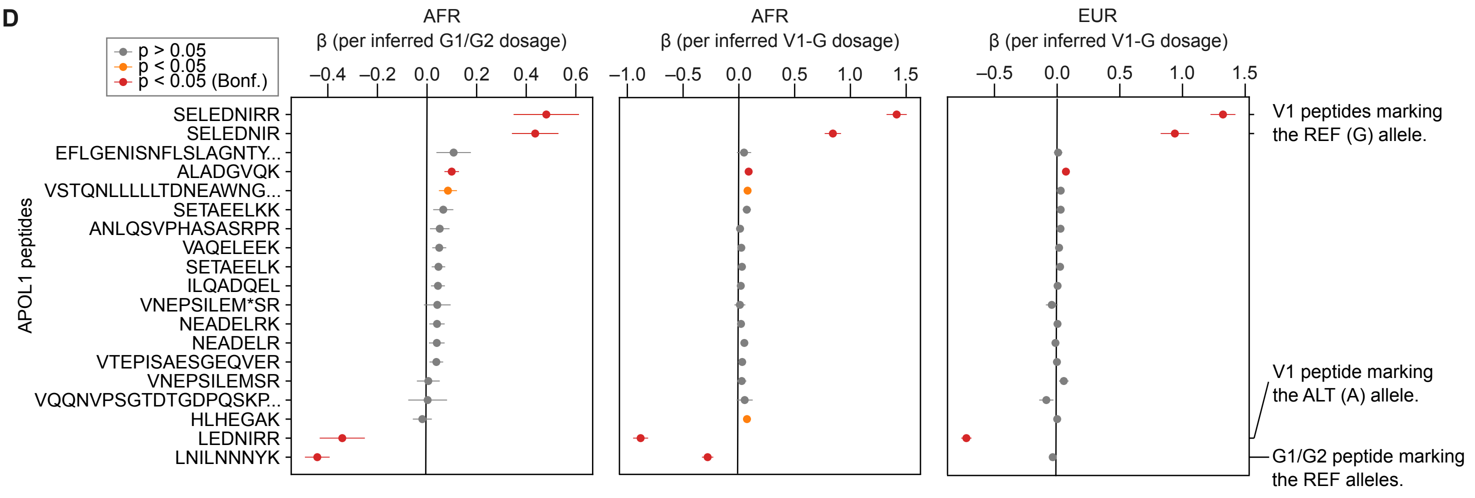

**Figure S4**

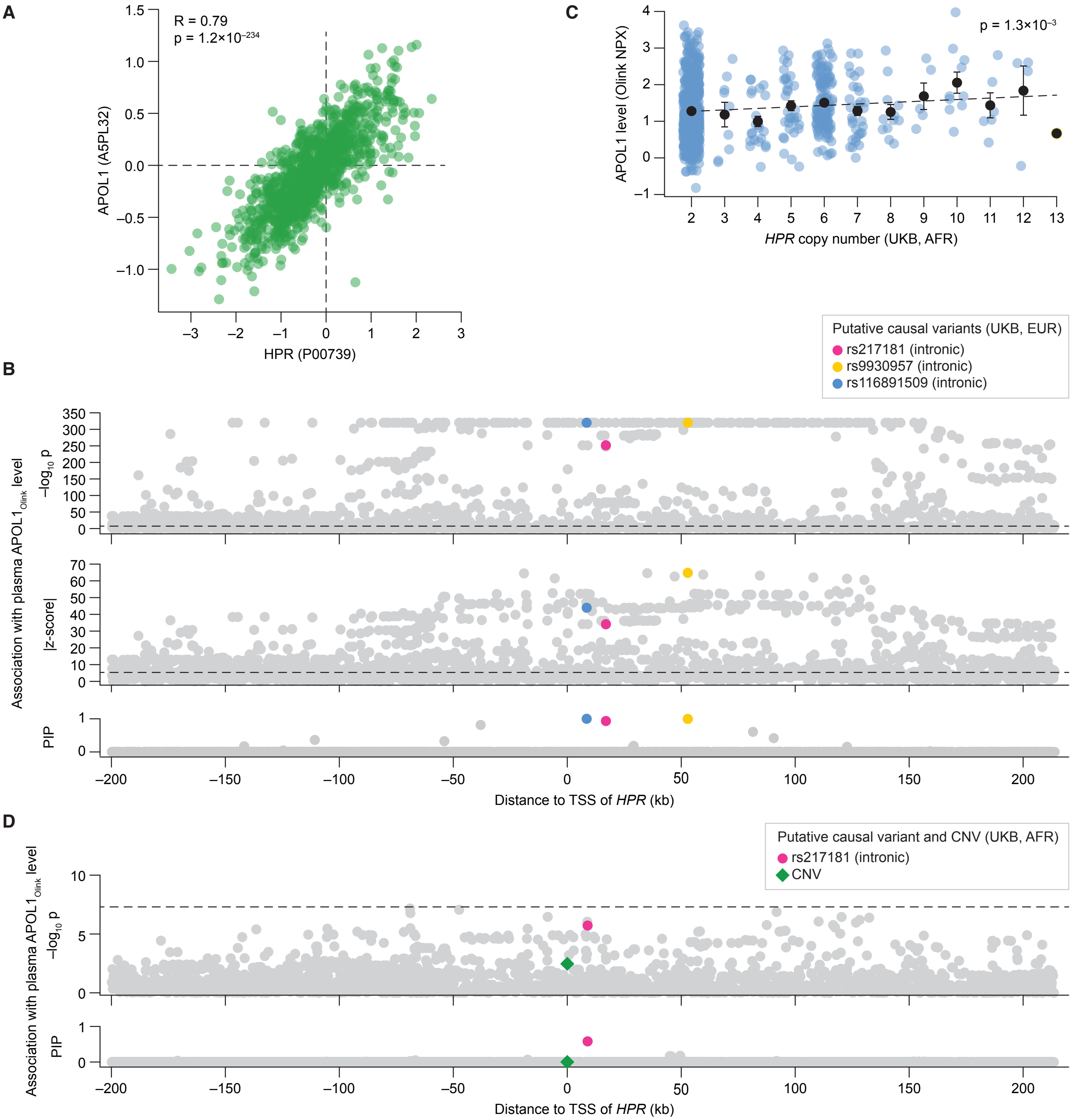


**Supplementary Figure 4. HPR cis-variants, but not copy number variation, are associated with plasma APOL1$_{Olink}$ levels. (A)** Scatter plot comparing plasma levels of HPR (x-axis) and APOL1 (y-axis) measured by mass spectrometry in the same individuals from Health ABC. **(B)** Manhattan plot showing the association of *HPR cis*-variants with plasma APOL1$_{Olink}$ levels in the UK Biobank population of European descent. Each dot represents a variant. The x-axis shows the variant's chromosomal position relative to the transcriptional start site (TSS) of *HPR*. The y-axis shows the $-\log_{10}$ of the association p-value (capped at 320; row 1), the absolute value of the association z-score (row 2) and the statistical fine-mapping posterior inclusion probability (PIP; row 3). Putative causal variants identified through fine-mapping are highlighted. Horizontal dashed lines represent the genome-wide significance threshold ($p = 5\times10^{-8}$). **(C)** Scatter plot comparing estimated *HPR* copy number (x-axis) and plasma APOL1$_{Olink}$ levels (y-axis) in the UK Biobank population of African descent. Each dot represents an individual. Within each copy number group, dots were jittered along the x-axis for visualization purposes. Black dots and corresponding error bars indicate the mean and standard error of the mean of each copy number group. The horizontal dashed line represents a linear regression fit. **(D)** Manhattan plot showing the association of *HPR cis*-variants and copy number variation (CNV) with plasma APOL1$_{Olink}$ levels in the UK Biobank population of African descent. Each dot represents a variant. The x-axis shows the variant's chromosomal position relative to the TSS of *HPR*, except for the CNV, which is plotted at the TSS directly. The y-axis shows the $-\log_{10}$ of the association p-value (row 1) and the statistical fine-mapping PIP (row 2). Putative causal variants identified through fine-mapping and the CNV are highlighted. Horizontal dashed line represents the genome-wide significance threshold ($p = 5\times10^{-8}$).

**Figure S5**

A

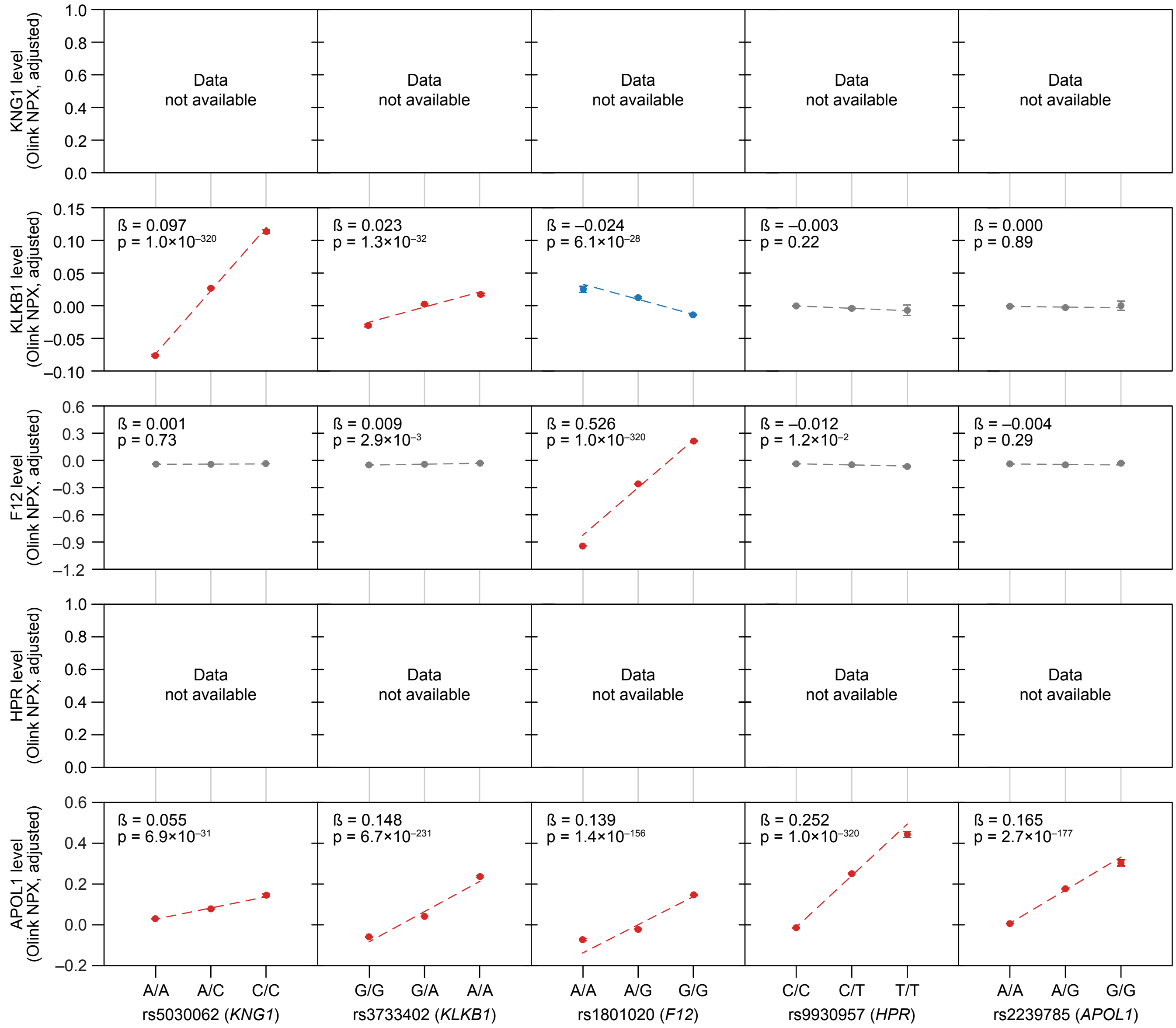


B

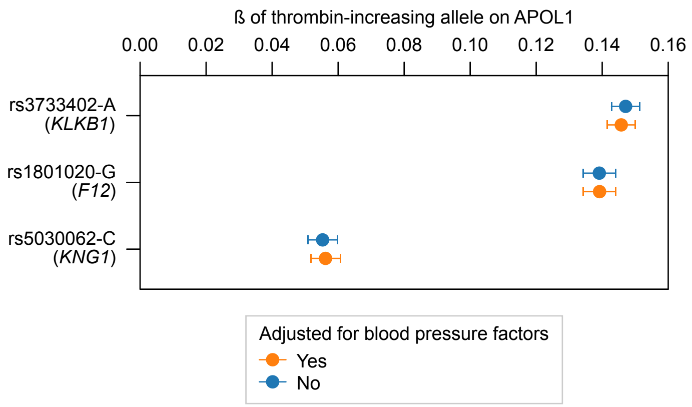


**Supplementary Figure 5. Comparison of pQTL effects suggests a positive correlation between the activity of the kallikrein-kinin pathway and plasma APOL1$_{Olink}$ levels. (A)** Dot plots showing the effects of KNG1, *KLKB1*, *F12*, *HPR*, and *APOL1* variants on covariate-adjusted plasma levels of KLKB1, F12, and APOL1$_{Olink}$ in individuals with European ancestry from the UK Biobank (Methods). Data for plasma KNG1 and HPR were not available as these proteins were not included in the Olink Explore 3072 platform used for the measurements. The mean protein level (y-axis) is shown for each variant genotype (x-axis). Error bars indicate standard errors of the mean. Associations with a significant positive or negative effect ($p < 5\times10^{-8}$) are colored in red and blue, respectively. **(B)** Forest plot showing the association of thrombin-increasing alleles of *KLKB1* (rs3733402-A), *F12* (rs1801020-G) and *KNG1* (rs5030062-C) with plasma APOL1$_{Olink}$ levels, both with and without adjustment for blood pressure-related factors (Methods). Effect sizes are reported as β on covariate adjusted APOL1$_{Olink}$ level (Methods). Error bars indicate standard errors of the mean.

**Figure S6**

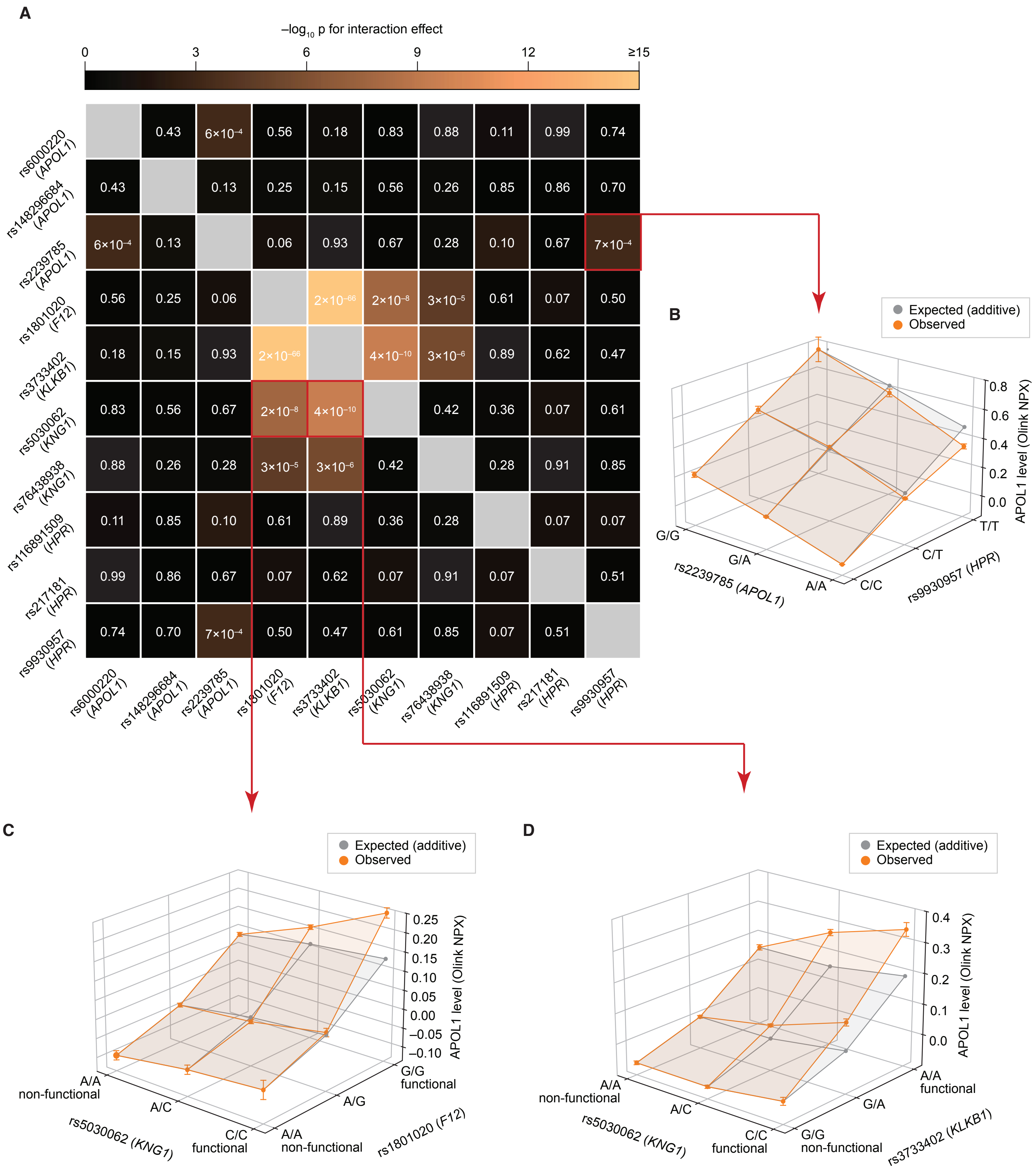


**Supplementary Figure 6. Variants in *KNG1*x*F12*, *KNG1*x*KLKB1*, and *HPR*x*APOL1* genetically interact to influence plasma APOL1$_{Olink}$ levels. (A)** Heatmap showing the significance of pairwise genetic interactions between all major putative causal pQTLs for plasma APOL1$_{Olink}$ levels. Cell color indicates the $-\log_{10}$ of the interaction p-value (Methods). **(B–D)** Examples of interacting variant pairs, in addition to those displayed in Fig. 6. In each plot, the x- and y-axes show variant genotypes, and the z-axis shows observed (orange) and expected, assuming additivity (grey), mean plasma APOL1$_{Olink}$ levels for each genotype combination. Error bars indicate the standard error of the mean.

**Figure S7**

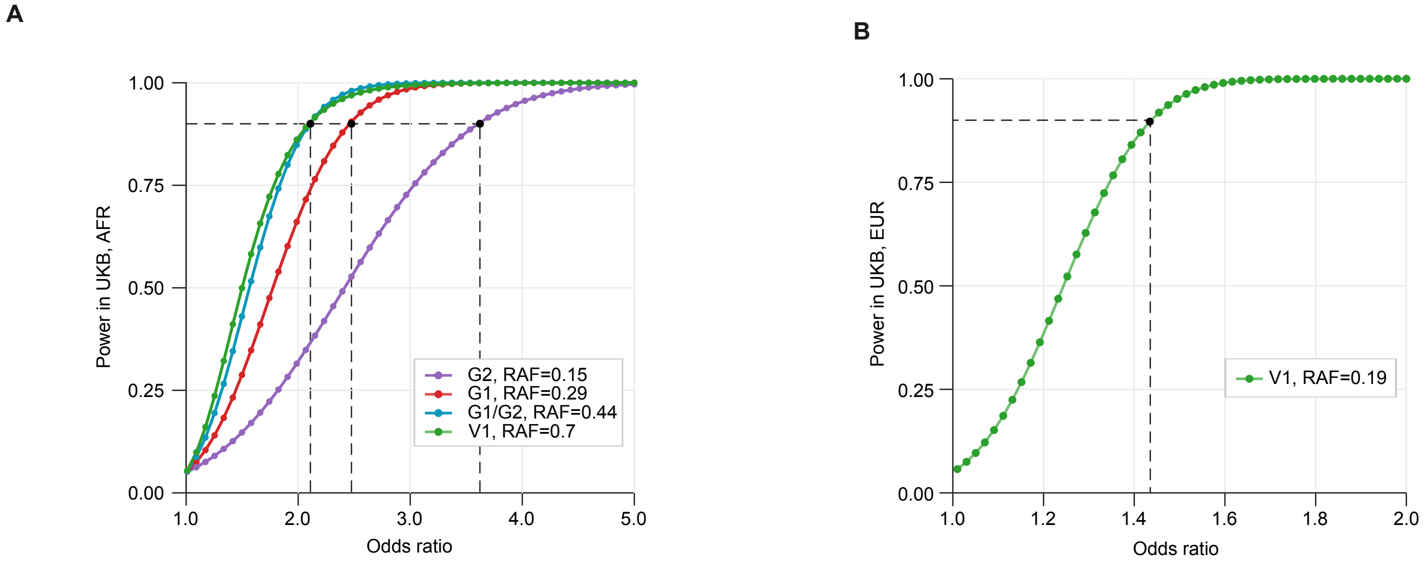


**Supplementary Figure 7. Power analysis suggests limited ability to detect the effect of G1/G2 and rs2239785 (V1) variants on end-stage kidney disease (ESRD) in the UK Biobank. (A–B)** Power to detect the variant effect at a significance level of α = 0.05 was calculated for a range of expected odds ratios, given the size of the UK Biobank population with (A) African (UKB, AFR) and (B) European ancestry (UKB, EUR) and the prevalence of ESRD and risk allele frequency (RAF) in that population (Methods). Power (y-axis) is shown as a function of the odds ratio (x-axis). Dashed lines indicate the minimal odds ratio detectable at 90% power.

**Figure S8**

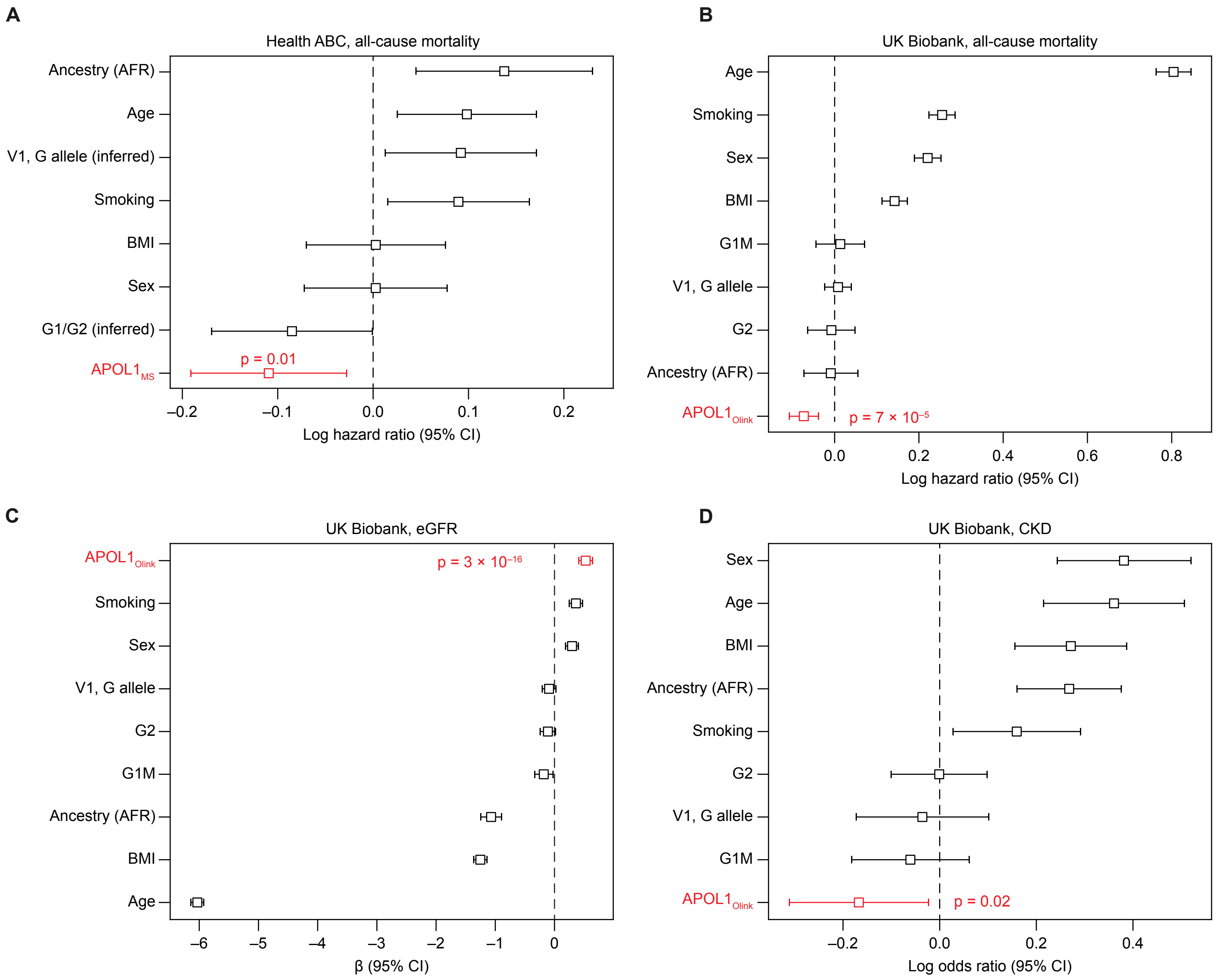


**Supplementary Figure 7. Higher plasma APOL1 levels are associated with a reduced risk of all-cause mortality and kidney disease. (A–B)** Association between plasma APOL1 levels and all-cause mortality in the Health ABC cohort (A) and the UK Biobank (B). A Cox proportional hazards model was used to assess the effect of baseline plasma $APOL1_{MS}$ (A) and $APOL1_{Olink}$ levels (B) on future mortality events, adjusting for age at recruitment, sex (0 = female, 1 = male), genetic ancestry (0 = European, 1 = African), smoking status (0 = never, 1 = ever) and body mass index (BMI). The forest plots display the log hazard ratio (HR) and the 95% confidence interval for each variable. **(C)** Association between plasma $APOL1_{Olink}$ levels and estimated glomerular filtration rate (eGFR) in the UK Biobank. A linear regression model was used to assess the relationship between plasma $APOL1_{Olink}$ levels and eGFR, adjusting for age at recruitment, sex, genetic ancestry, smoking status and BMI. The forest plot displays the regression coefficients for each variable. eGFR was calculated using the 2021 CKD-EPI creatinine formula (Inker *et al.*, 2021). **(D)** Association between plasma $APOL1_{Olink}$ levels and CKD diagnosis in the UK Biobank. A logistic regression model was used to assess the relationship between plasma $APOL1_{Olink}$ levels and CKD, adjusting for age at recruitment, sex, genetic ancestry, smoking status and BMI. The forest plot displays the regression coefficients for each variable. CKD diagnosis was determined using ICD-10 code N18.
